# Cryo-EM structure of SARS-CoV-2 postfusion spike in membrane

**DOI:** 10.1101/2022.12.05.519151

**Authors:** Wei Shi, Yongfei Cai, Haisun Zhu, Hanqin Peng, Jewel Voyer, Sophia Rits-Volloch, Hong Cao, Megan L. Mayer, Kangkang Song, Chen Xu, Jianming Lu, Jun Zhang, Bing Chen

## Abstract

Entry of severe acute respiratory syndrome coronavirus 2 (SARS-CoV-2) into host cells depends on refolding of the virus-encoded spike protein from a prefusion conformation, metastable after cleavage, to a lower energy, stable postfusion conformation. This transition overcomes kinetic barriers for fusion of viral and target cell membranes. We report here a cryo-EM structure of the intact postfusion spike in a lipid bilayer that represents single-membrane product of the fusion reaction. The structure provides structural definition of the functionally critical membrane-interacting segments, including the fusion peptide and transmembrane anchor. The internal fusion peptide forms a hairpin-like wedge that spans almost the entire lipid bilayer and the transmembrane segment wraps around the fusion peptide at the last stage of membrane fusion. These results advance our understanding of the spike protein in a membrane environment and may guide development of intervention strategies.

## Introduction

The COVID-19 (coronavirus disease 2019) pandemic, caused by severe acute respiratory syndrome coronavirus 2 (SARS-CoV-2), has a death toll in millions and devastating socioeconomic impacts worldwide. SARS-CoV-2 is an enveloped positive-stranded RNA virus that enters a host cell after fusion of the viral and cell membranes. Although membrane fusion is energetically favorable, there are high kinetic barriers when two membranes approach each other, primarily because of repulsive hydration forces (*1*, *2*). Free energy, required for viral membrane fusion to overcome the kinetic barriers, comes from refolding of the virus-encoded fusion protein from a prefusion conformation, which is metastable after proteolytic cleavage to a lower energy, stable postfusion state (*3*–*5*). The fusion protein of SARS-CoV-2 is its spike (S) protein, which is a type I, heavily glycosylated membrane protein with a transmembrane (TM) segment embedded in the viral membrane, and another membrane-interacting region, known as “fusion peptide (FP)”, which can insert into the target cell membrane (*6*). Like other class I viral fusion proteins including HIV-1 envelope glycoprotein, influenza hemagglutinin and Ebola glycoprotein (*3*, *4*), the S protein is synthesized as a single-chain precursor, trimerized and subsequently cleaved by a furin-like protease from the infected host cell into two fragments: the receptor-binding fragment S1 and the fusion fragment S2 (*7*). To initiate the next round infection, S protein binds to the receptor angiotensin converting enzyme 2 (ACE2) on the surface of a new host cell and is further cleaved at a second site in S2 (S2’ site) by a host serine protease TMPRSS2 or an endosomal cysteine protease cathepsin L (*6*, *8*, *9*). It then undergoes large conformational changes to insert the FP into the target cell membrane and then refold into a hairpin-like postfusion structure, placing the TM and FP at the same end of the molecule, thereby dragging two membranes close together to complete fusion (*10*, *11*).

Soon after the release of the first SARS-CoV-2 genomic sequence (*12*), the structures of the S protein fragments, such as its ectodomain stabilized in the prefusion conformation (*13*, *14*), the receptor binding domain (RBD) in complex with ACE2 (*15*–*18*) and fragments of S2 in the postfusion conformation (*19*), were reported. In the prefusion ectodomain structure, S1 folds into four different domains - N-terminal domain (NTD), RBD, C-terminal domain 1 (CTD-1) and C-terminal domain 2 (CTD-2), and they wrap around the prefusion conformation of S2. The structures of the purified full-length S protein in both the prefusion and postfusion conformations (*20*, *21*), as well as those of the S proteins present on chemically inactivated SARS-CoV-2 virions (*22*–*25*), were subsequently determined, revealing additional structural details. The fusion fragment S2 contains several segments that play important structural and functional roles, including the FP, heptad repeat 1 (HR1), central helix (CH), connector domain (CD), heptad repeat 2 (HR2), TM and cytoplasmic tail (CT) (Fig. S1). In the postfusion trimer of the S2 ectodomain, HR1 and CH form a central three-stranded coiled coil, ~180Å long (*20*). Part of HR2 adopts a helical conformation and packs against the groove of the HR1 coiled-coil to form a six-helix bundle and stabilize the hairpin-like postfusion structure. It is consistent with a membrane fusion model, in which HR1 undergoes a “jack-knife” transition to insert the FP into the target cell membrane and HR2 folds back to bring the FP and TM segments close together (*7*), in turn causing the two membranes to fuse into a single lipid bilayer. In all previous structures, however, the regions of the S protein near the viral membrane, while structurally and functionally critical (*26*–*30*), are either not present or disordered.

Among the membrane-interacting regions of coronavirus spike proteins, the FP is one of the most crucial structural elements for membrane fusion, but its exact location has been disputed (*31*). Three membranotropic regions in the SARS-CoV (severe acute respiratory syndrome coronavirus) S2 have been suggested as putative fusion peptides (Fig. S1), including a potentially glycosylated segment upstream of the S2’ cleavage site (residues 770-788, named the N-terminal FP (n-FP); ref (*32*, *33*)); the segment immediately downstream of the S2’ cleavage site (residues 798-816 or 798-835, widely accepted as the *“bona fide”* FP (b-FP); ref (*34*, *35*)); and the segment immediately upstream of HR1 (residues 858-886, also known as the internal FP (i-FP); *ref*(*29*, *33*, *36*)). Early studies supported the assignment of the i-FP because it had strong membrane-perturbing capacities and mutations in this region led to inhibition of the S-mediated cell-cell fusion (*29*). Subsequent studies have shown that the b-FP, highly conserved among coronaviruses, has even a stronger calcium-dependent membrane ordering activity -- a main characteristic of a viral FP when studied as a short peptide in solution -- than the i-FP (*34*, *35*). Recently, several broadly neutralizing antibodies have been identified, recognizing the b-FP region of spike proteins from all known human-infecting coronaviruses, and the fusion peptide has therefore been suggested as a potential target for developing universal coronavirus vaccines (*37*, *38*). A fourth hydrophobic region (residues 1190-1203, also called the pretransmembrane domain (pre-TM) or aromatic domain) adjacent to the TM domain has been shown to be important in SARS-CoV fusion, and may act in concert with the FP to support membrane fusion (*39*, *40*).

To visualize the membrane interacting regions of postfusion SARS-CoV-2 spike protein, we have reconstituted the full-length protein in lipid-based nanodiscs, induced the conformational changes using soluble ACE2 to prepare a sample of the postfusion spike in lipid bilayer, and determined its structure by cryogenic electron microscopy (cryo-EM) to reveal structural details critical for a full understanding of the SARS-CoV-2 entry.

## Results

### Preparation of the postfusion S2 trimer in lipid nanodiscs

We reconstituted the purified, intact protein derived from the early variant G614 (B.1) in lipid nanodiscs using a circularized membrane scaffold protein (MSP; ref (*41*, *42*)). The reconstituted sample resolved by gel-filtration chromatography into three major peaks. SDS-PAGE analysis showed that peak I contained mainly the cleaved S protein in nanodiscs, peak II, empty nanodiscs without the S protein, and peak III, the free MSP (Fig. 1A). Negative stain EM confirmed the identity of peak I as the nanodisc-associated prefusion S trimer, showing some flexibility between the ectodomain and the TM region. We induced the conformational changes of the prefusion S trimer by incubating the reconstitution reaction with soluble ACE2. The ACE2-treated sample also resolved into three peaks by gel-filtration chromatography. Peak I contained primarily the dissociated S2 in nanodiscs, peak II, empty nanodiscs co-eluting with dissociated S1 in complex with ACE2, and peak III, unbound ACE2 co-eluting with free MSP (Fig. 1B). Negative stain EM images of the peak I fraction showed a very rigid, postfusion S2 trimer projecting from the nanodisc. These results demonstrate that ACE2 binding is sufficient to trigger the prefusion-to-postfusion conformational transition of the purified S trimer even in the context of the restricted bilayer membrane in a nanodisc.

**Figure 1.**
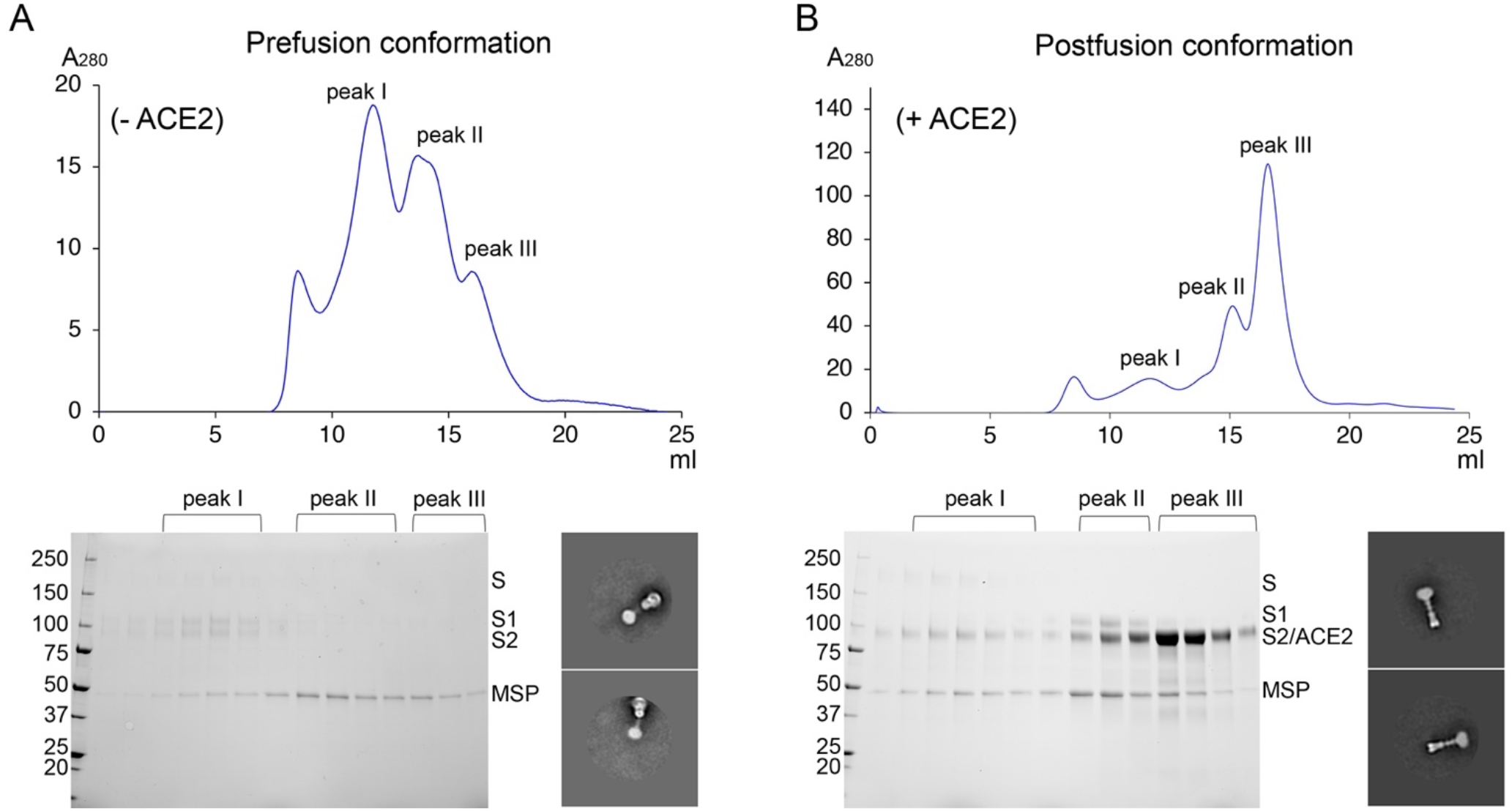
Preparation of the membrane-bound SARS-CoV-2 spike protein in lipid nanodiscs. (**A**) The purified full-length SARS-CoV-2 S protein was reconstituted in lipid nanodiscs and resolved by gel-filtration chromatography on a Superose 6 column. Three major peaks (not at the void volume) are peak I containing the prefusion S trimer in nanodiscs, peak II containing empty nanodiscs, and peak III containing free membrane scaffold protein (MSP), as analyzed by SDS-PAGE. Representative 2D averages by negative stain EM of the peak I fractions are also shown. The box size of 2D averages is ~880Å. (**B**) The purified full-length SARS-CoV-2 S protein was first reconstituted in nanodiscs, incubated with soluble ACE2 and then resolved by gel-filtration chromatography on a Superose 6 column. Three non void-volume peaks are peak I containing the postfusion S2 trimer in nanodiscs, peak II containing empty nanodiscs and dissociated S1 in complex with ACE2, and peak III containing unbound ACE2 and free MSP, as analyzed by SDS-PAGE. Representative 2D averages by negative stain EM of the peak I fractions are also shown. The box size of 2D averages is ~880Å.

### Cryo-EM structures of the membrane-bound postfusion S2 trimer

We determined by cryo-EM the structure of postfusion S2 trimer in nanodiscs, prepared as just described. The terminology for various segments of the S2 polypeptide chain are depicted in Fig. 2A. We recorded cryo-EM images on a Titan Krios electron microscope equipped with a Gatan K3 direct electron detector and used cryoSPARC (*43*) for particle picking, two-dimensional (2D) classification, three-dimensional (3D) classification and refinement (Fig. S2). One major class was obtained from 3D classification and refined to 2.9Å resolution (Fig. S2-S4; Fig. 2B; Table S1). To improve the local resolution near the nanodisc, we performed additional masked local refinement, leading to a 3.3 Å map covering the transmembrane and cytoplasmic tail regions.

**Figure 2.**
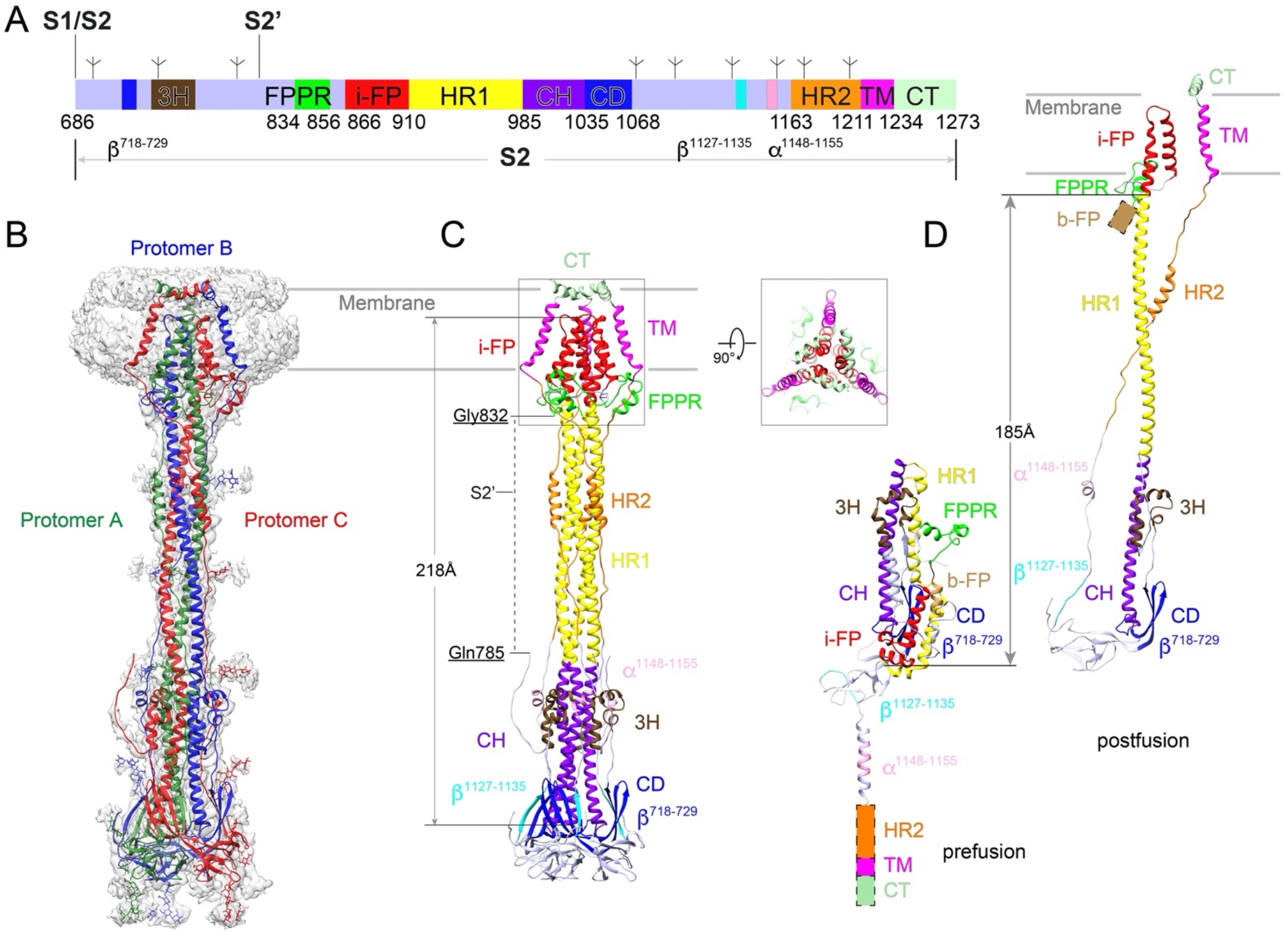
Cryo-EM structure of the SARS-CoV-2 S2 in the postfusion conformation. (**A**) Schematic representation of the SARS-CoV-2 S2 fragment. Various segments include: S1/S2, S1/S2 cleavage site; S2’, S2’ cleavage site; β^718-729^, a β-strand formed by residues 718-729 in the S1/S2-S2’ fragment; 3H, three-helix segment; FPPR, fusion peptide proximal region; i-FP, the internal fusion peptide; HR1, heptad repeat 1; CH, central helix region; CD, connector domain; β^1127-1135^, a β-strand formed by residues 1127-1135; a^1148-1155^, an a-helix formed by residues 1148-1155; HR2, heptad repeat 2; TM, transmembrane anchor; CT, cytoplasmic tail; and tree-like symbols for glycans. (**B**) The structure of the S2 trimer fits into a 2.9Å density map. Three protomers (A, B, C) are colored in green, blue and red, respectively. (**C**) Overall structure of the S2 trimer in the postfusion conformation shown in ribbon diagram. Various structural components in the color scheme shown in (A) include 3H, FPPR, i-FP, HR1, CH, CD, HR2, TM and CT. The S2’ cleavage site is in a disordered segment between Gln785 and Gly832, indicated by a dashed line. A top view of the transmembrane region in the nanodisc is also shown. (**D**) One protomer each of the S2 trimer from the prefusion and the postfusion conformations superposed by the invariant CH and CD region. The locations of HR2, TM and CT disordered in the prefusion structure, as well as the b-FP in the postfusion structure are indicated by colored rectangles. The N-terminal end of HR1, connected directly to the i-FP, translocates by ~185Å during the prefusion-to-postfusion transition.

The overall structure of the postfusion S2 ectodomain in a nanodisc is nearly identical to that of the protein solubilized in detergent (*20*) and that of SARS-CoV (*44*). As reported previously, the core structure is a long central three-stranded coiled coil made up by HR1 and CH (Fig. 2C). A three-stranded ß sheet formed by a β hairpin from the connector domain (CD), and a segment (residues 718-729; β^718-729^) from the S1/S2-S2’ fragment, created by proteolytic cleavages, wraps around the C-terminal end of the CH coiled-coil. This connector β sheet and CH form the invariant structure between the prefusion and postfusion structures. Another segment (residues 737-769) in the S1/S2-S2’ fragment folds into three consecutive a-helices (3H), locked by two disulfide bonds, and also tightly packing against the groove of the CH coiled-coil. In our new structure, there is additional density for the segment immediately downstream 3H, which bends back towards the TM region and attaches to the surface of the postfusion structure, becoming disordered after Gln785. The S2’ site near Arg815, presumably cleaved in this conformational state, remains invisible. In the C-terminal half of S2, a β strand formed by residues 1127-1135 (β^1127-1135^) upstream of HR2 from another protomer joins the connector β sheet to expand it into four strands, projecting HR2 towards the TM region and probably initiating the HR2 folding back. Furthermore, a two-turn helix formed by residues 1148-1155 (a^1148-1155^) wedges between two neighboring 3Hs and a longer helix of HR2 makes up the six-helix bundle with the HR1 coiled-coil, together reenforcing the very rigid postfusion structure.

The membrane-interacting segments of S2, missing in all previous postfusion structures of any class I viral fusion proteins, are well resolved in our new structure. There are nine membrane-spanning helices (three per protomer) in the nanodisc region, which were fully resolved in the refined maps. The i-FP region immediately upstream of HR1 forms a continuous a-helix extending the central coiled-coil to ~218Å long and well into the lipid bilayer (Fig. 2C), possibly accounting for the rigidity of the entire postfusion structure including the transmembrane region. This first i-

FP membrane-spanning helix is followed by a sharp U-turn within the lipid bilayer, and another helix that spans through the membrane once again and sends the adjacent FPPR back to the ectodomain side of the membrane. A total of six transmembrane helices of the i-FP from three protomers pack tightly together to form a blunted cone shape (Fig. 2C), probably required for effectively penetrating the target cell membrane. The third membrane-spanning helix is the TM segment, which tilts relative to the plane of the membrane, gently wrapping around the blunted cone. Part of the following CT appears to be embedded horizontally in the cytosolic headgroup region of the lipid bilayer, and three copies of it form a triangle that caps the tip of the transmembrane cone (Fig. 2C). Comparison of the prefusion and postfusion structures indicates that formation of the long postfusion central helix translocates by 185 Å the i-FP, which reconfigures to become a hairpin-like wedge inserted into the target membrane (Fig. 2D). The i-FP not only inserts into the target membrane but also becomes a docking site for the TM helix in the final stages of membrane merger.

### Structural definition of the SARS-CoV-2 fusion peptide and transmembrane segment

The local refined maps with sidechain density for many residues in the transmembrane region allowed unambiguous modeling (Fig. S4). It is clear that both the n-FP (residues 788-806; Fig. S5) and b-FP (residues 816-834; Fig. S5) are disordered but located outside the membrane (Fig. 2C), and thus they do not function as the real fusion peptide. It is the i-FP that forms a hairpin-like wedge in a trimeric cone-shaped assembly to span almost the entire lipid bilayer (Fig. 3A). As expected, the membrane-inserted region of the wedge is largely hydrophobic with Trp885 and Phe888 at the blunted tip while there are several charged residues (Asp867, Glu868 and Arg905) at the base of the wedge located in the headgroup region of lipid bilayer. Thus, total 44 residues (Asp866 - Ile909) insert into membrane and make up the functional fusion peptide. Interestingly, Pro897 introduces a kink in the long central helix, making the region near the tip of the fusion wedge deviated from a canonical coiled-coil structure and also allowing the tight packing of the returning helix (residues 870-882) through an ^876^AXXXG^880^ motif to form the blunted cone shape (Fig. 3A).

**Figure 3.**
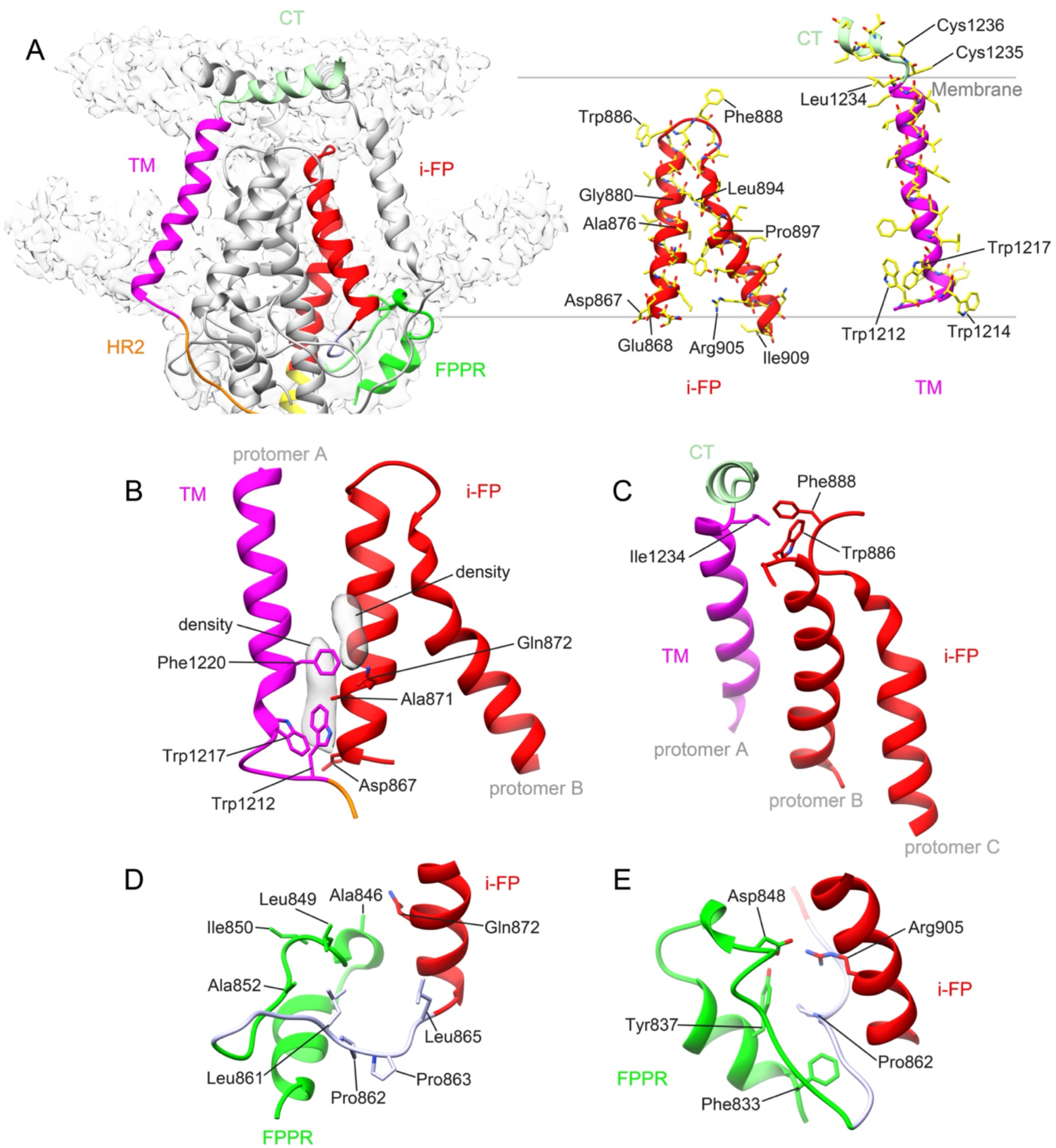
Structural details of the postfusion SARS-CoV-2 spike. (**A**) Left, the transmembrane region of the postfusion S2 fits in the EM density with one protomer colored: HR1 in yellow, FPPR in green, i-FP in red, HR2 in orange, TM in magenta and CT in light green. Right, the atomic models for the i-FP and TM segment with key residues indicated. (**B**) Interactions between the TM and i-FP near the ectodomain side of the membrane with key interface residues shown in stick model. The elongated density potentially for ordered phospholipids are shown in gray. (**C**) Interactions between the TM and i-FP near the CT side of the membrane with key interface residues shown in stick model. (**D**) Attachment of the FPPR and the segment of residues 857-866 to the lipid bilayer with the hydrophobic residues facing towards the hydrophobic core of the membrane highlighted in stick model. (**E**) Interactions between the FPPR and i-FP on the ectodomain side of the membrane with key interface residues shown in stick model.

For the transmembrane anchor, the TM helix begins at Tyr1215 and ends at Leu1234, and Trp1212 and Trp1217 from the pre-TM are embedded entirely in the lipid bilayer, indicating that the pre-TM is part of the TM domain, not a separate structural element. In addition, Cys1235 and Cys1236 are clearly not part of the TM helix and mark the beginning of the CT (Fig. 3A). These two cysteine residues, possibly palmitoylated in the virus (*45*, *46*), are also very close to Cys1247 and Cys1248 from the CT of a neighboring protomer, although formation of disulfide bonds between them on the cytosolic side is unlikely.

### Interactions between the fusion peptide and other membrane-interacting segments

It is postulated that the FP of a class I viral fusion protein may interact with its TM domain in the final stage of the fusion process, but there has been no direct structural evidence for this interaction in an intact protein in the context of membrane. The structure reported here shows an intimate FP-TM interaction in the SARS-CoV-2 postfusion spike. The TM segment packs against the fusion peptide from another protomer with two major contacts. First, Trp1212, Trp1217 and Phe1220 near the ectodomain side pack against the base of the fusion wedge; the contact may also involve some phospholipids, as suggested by strong elongated density in the vicinity, aligned perpendicular to the membrane plane (Fig. 3B). Second, Ile1234 near the CT side interacts with Trp886 and Phe888 from two different protomers at the tip of the fusion wedge (Fig. 3C). Three copies of the CT fragment come together to cap the tip of the fusion wedge.

The segment (residues 857-866) immediately upstream of the i-FP lies in the headgroup region of the lipid bilayer with several hydrophobic residues pointing towards the hydrophobic core of the membrane. This configuration probably helps orient the adjacent disulfide-locked FPPR (residues 835-856) in a way with its C-terminal half embedded in the membrane as well (Fig. 3D). The N-terminal half of the FPPR, directly connected to the disordered b-FP, projects away from the membrane. In addition, the FPPR packs against the i-FP wedge and Asp848 from the FPPR forms a salt bridge with Arg905 of the i-FP. Thus, the FPPR, which facilitates clamping down of the RBD in the prefusion conformation (*20*), may also contribute to anchoring the membrane-interacting regions of the postfusion S2 structure.

### Effect of mutations in the membrane-interacting segments on membrane fusion

To test the functional roles of the membrane-interacting structural elements, we have assessed impacts of several structure-guided mutations in the b-FP, i-FP, FPPR and CT regions. As illustrated in Fig. 4A, we substituted multiple hydrophobic residues in the b-FP, which would in principle be critical for membrane interaction, with charged residues (Mut-1); we mutated the key residues that participate in the FPPR interaction with the i-FP (Mut-2); we introduced multiple charged residues in the i-FP (Mut-3 and Mut-4) and mutated its AXXXG motif (Mut-5 and Mut-6); and we changed the two cysteine residues that mark the beginning of the CT (Mut-7).

**Figure 4.**
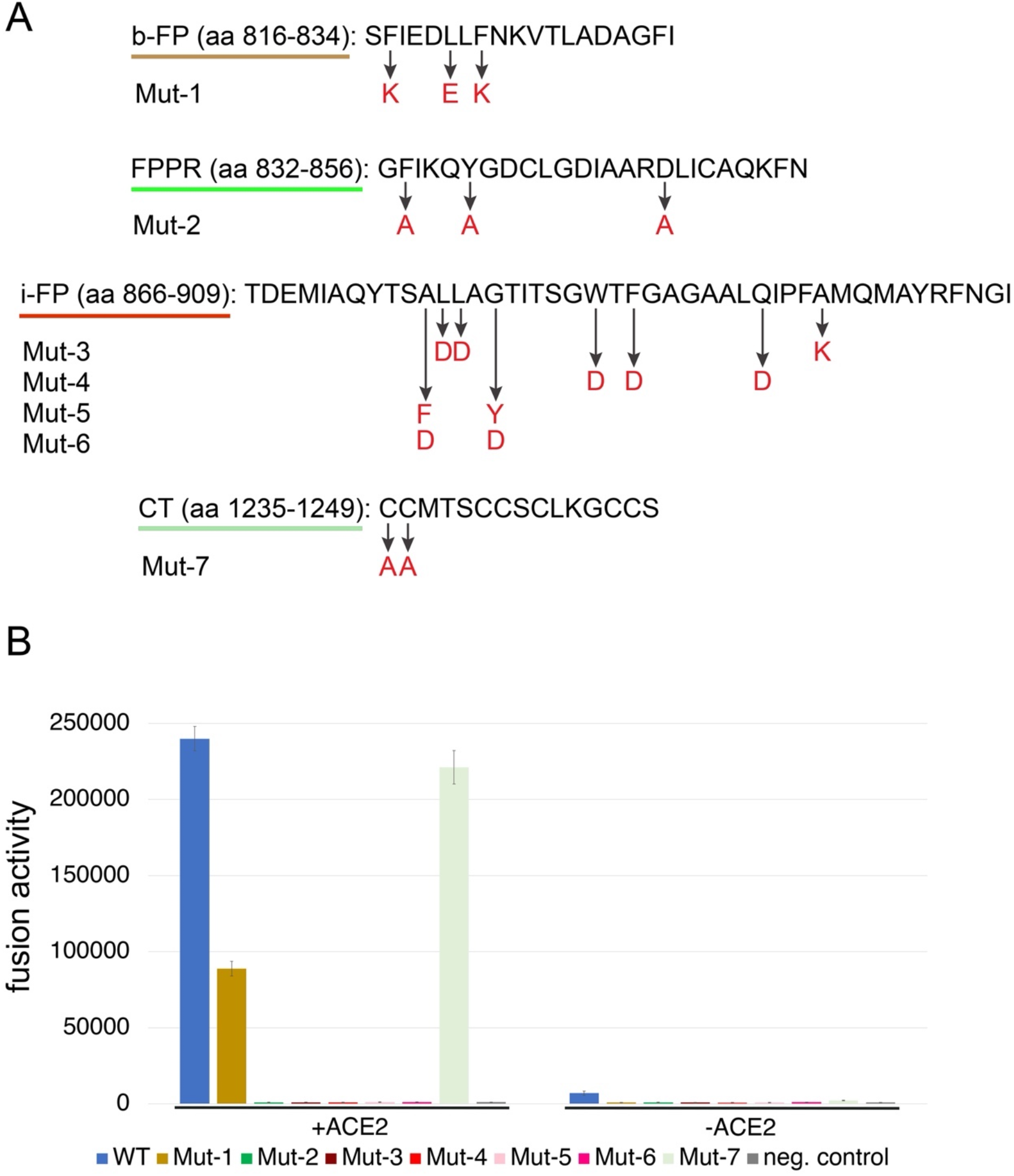
Effect of mutations in the membrane-interacting segments on membrane fusion. (**A**) Structure-guided design of mutations in the b-FP, FPPR, i-FP and CT regions. The wildtype sequences are shown in black and the mutated residues in red. (**B**) Cell-cell fusion mediated by the G614 S and seven designed mutants. HEK293T cells transfected with the full-length S protein expression plasmids were fused with ACE2-expressing cells. Cell-cell fusion led to reconstitution of a and ω fragments of β-galactosidase to form an active enzyme, and the fusion activity was then quantified by a chemiluminescent assay. No ACE2 were negative controls. The experiment was repeated four times with similar results. Error bars, mean ± s.d.

When transiently transfected in 293T cells, these mutants expressed the same level of S, but the extent of cleavage between S1 and S2 was substantially reduced for Mut-3, Mut-4, Mut-5 and Mut-6, all of which are in the i-FP (Fig. S6A). Like fusion peptides and fusion loops of many other viral fusion proteins (*47*–*49*), the i-FP is buried in the prefusion conformation, in the present case by a segment near the N-terminus of S2 from another promoter (*13*, *14*, *20*). When detected by monoclonal antibodies using flow cytometry (Fig. S6B), the recognition patterns by RBD-specific antibodies (C63C8, G32B6, C63C7, REGN10933 and REGN10987; ref (*50*, *51*)), NTD-specific antibodies (C12C9, 4A8 and C81D6; ref (*50*, *52*)), S2-specific antibodies (C163E6 and 3H3; ref (*50*, *52*)) and ACE2 for Mut-1, Mut-2 and Mut-7 were very similar to that of the wildtype S, except that the overall binding levels were substantially lower for Mut-1, suggesting that the mutations had some impact on protein secretion. The secretion level of Mut-3 was very low, as judged by binding to all antibodies and ACE2, indicating that this mutant protein might not be folded correctly. The patterns for binding to the antibodies and ACE2 were similar among Mut-4, Mut-5 and Mut-6, but they bound more weakly to RBD-specific antibodies and the neutralizing NTD-specific antibody, 4A8, while much more strongly to S2-specific antibodies and to the nonneutralizing NTD-specific antibody, C81D6, than did the wildtype S, consistent with large structural rearrangements caused by the mutations in these proteins. These results suggest that the i-FP, largely buried in the prefusion structure, does not tolerate most mutations, even in the prefusion conformation, consistent with its buried position. In a S-mediated cell-cell fusion assay (*20*), Mut-1 showed reduced fusion activity while Mut-7 behaved like a wildtype spike; all other mutants showed no detectable fusion activity (Fig. 4B). Thus, as our structure shows, the b-FP is not the fusion peptide and does not interact with membrane, as introducing multiple charged residues does not abolish membrane fusion. Moreover, the two cysteine residues in the CT close to the TMD, which have the potential to form interchain disulfide bonds, are dispensable for membrane fusion. It is more complicated to interpret the data for Mut-3, Mut-4, Mut-5 and Mut-6 since their mutations appeared to have substantial impact on the prefusion structures of the mutant proteins. Therefore, complete lack of membrane fusion activity by these mutants may be attributed to effects on the structural role of the i-FP in the prefusion state, as well as the functional role as the fusion peptide in the postfusion conformation. Nevertheless, these results further confirm that the i-FP is a key functional structural element in the SARS-CoV-2 spike.

By molecular modeling using AlphaFold (*53*), most mutations in the SARS-CoV-2 variants of concern are at surface-exposed residues in the postfusion structure except for the Omicron subvariants, in which some residues such as L981F may help stabilize the structure while others such as N969K may destabilize it (Fig. S7).

## Discussion

Our structure of the SARS-CoV-2 postfusion S2, with membrane-associated elements interacting with a lipid bilayer, has resolved a long-standing contentious issue in the coronavirus field regarding the functional identity of the fusion peptide. Although both the n-FP and i-FP were suggested as candidates, the b-FP subsequently became widely accepted as *“bonafide”* because it was more conserved among coronaviruses and showed a stronger activity than the other two in perturbing membrane structure as a synthetic peptide (*34*, *35*). Our results show unambiguously that only the i-FP interacts with membrane. They underscore the risk of studying fusion peptides when taken out of the context of intact viral fusion proteins, because many hydrophobic peptides can induce structural changes of a lipid bilayer. For example, the FPPR, also identified as the “fusion peptide 2” for SARS-CoV in some studies (*35*), projects partway into the membrane headgroup region and could therefore modify the local membrane structure, but it does not insert into the hydrophobic part of the lipid bilayer. From our structure, neither the n-FP nor the b-FP inserts into the membrane in the postfusion structure and therefore neither is likely to function as a real “fusion peptide”. The conclusion is further supported by the evidence that introducing multiple charged residues in the b-FP does not abolish the membrane fusion activity of the fulllength S and by the presence of a conserved N-linked glycosylation site in the n-FP. We cannot rule out the possibility that the n-FP and b-FP may transiently interact with the membrane during the fusion process. Nevertheless, membrane insertion of the i-FP does not seem to depend on the n-FP or b-FP since it is probably directly coupled with the refolding of adjacent HR1 into the coiled-coil. Moreover, a fusion peptide may not have to be fully conserved in its amino acid sequence as long as it maintains the ability to insert correctly into the target membrane. Indeed, there is a four-residue deletion at the tip of the i-FP in some coronaviruses, including HCoV-OC43 and HCoV-229E (Fig. S5), but it is not likely to alter the overall shape of the fusion wedge and thus its conserved function.

The substantial lipid-bilayer interaction surface of the fusion wedge observed here is consistent with the stability of the extended intermediate, shown to persist for many minutes, even after release of S1 and its associated ACE2, when SARS-CoV-2 binds at the cell surface, unless the pH falls below neutral (*54*). Fusion peptides with less extensive hydrophobic contacts, such as those postulated for the influenza virus fusion peptide (*55*), would probably have shorter lifetimes.

Our structure also provides a more accurate structural definition of the TMD, as well as the direct evidence for interactions between the FP and TM in the merged membrane for a class I viral fusion protein. The previously designated “pre-TM” segment, now shown to be part of the TM, contributes to packing of the TM against the i-FP, explaining its functional role in membrane fusion. One study reported that a single residue insertion into the TM of SARS-CoV S led to a complete loss in membrane fusion (*56*), and the observation was difficult to interpret based on the presumptive trimeric TMD structure in the prefusion conformation (*57*). The insertion was between G1201 and F1202 of SARS-CoV, which correspond to G1219 and F1220 in SARS-CoV-2. Such an insertion would cause a rotation of the C-terminal portion of the TM helix possibly together with the CT in the postfusion structure, thereby disrupting the interactions between the i-FP and the TM and CT and blocking the last step of membrane fusion. These results, together with our new structure, highlight the functional importance of the interactions among the membrane-interacting elements within the lipid bilayer, which have not been recognized previously.

Finally, our structure suggests that the real fusion peptide of SARS-Cov-2 spike is unlikely a useful vaccine target since it is well protected and not accessible by antibodies in the prefusion conformation (*13*, *14*, *20*), nor is it exposed after inserting into the target cell membrane in the prehairpin intermediate state. The b-FP, targeted by several broadly neutralizing antibodies (*37*, *38*), may be conserved for other structural or functional reasons, which will require further investigation. Moreover, the interactions among the i-FP and FPPR, TM and CT shown in our structure, may provide some novel therapeutic targets for developing peptide-based or smallmolecule fusion inhibitors. In particular, mutations in the FPPR, designed to disrupt its interaction with the i-FP, completely abolish fusion activity, raising the possibility of identifying therapeutic candidates that target this interaction and block viral infection.

## Materials and Methods

### Methods

#### Expression constructs

The expression constructs for the full-length spike (S) protein (residue 1-1273) of SARS-CoV-2 (G614) and monomeric soluble ACE2 protein were described previously (*58*, *59*). To produce membrane scaffold protein (MSP), a modified gene encoding MSP2N2 (*41*, *42*) fused with a 7×His tag and a TEV site at N-terminal end and a sortase eSrt site and a 6×His tag at C-terminal end, was synthesized and cloned into the pET28a vector between *Nco* I and *Xho* I restriction sites to create the construct pET28a-lsMSP2N2 by GenScript (Piscataway, NJ). The expression constructs for the mutant spike proteins were generated by standard PCR methods and verified by DNA sequencing of the entire coding regions.

#### Expression and purification of recombinant proteins

A stable cell line was generated using HEK 293T cells for large-scale production of the full-length G614 S protein following the published protocol (*60*). Purification of the full-length S protein was carried out as previously described (*58*). Briefly, the stably transfected cells were grown in Expi293 expression medium (Thermo Fisher Scientific, Waltham, MA) containing 1% Pen Strep (Thermo Fisher Scientific) and 1.0 μg/ml puromycin (Thermo Fisher Scientific) to a density of ~4-5×10^6^/ml, then lysed in a lysis buffer containing Buffer A (100 mM Tris-HCl, pH 8.0, 150 mM NaCl, 1 mM EDTA) and 1% (w/v) n-dodecyl-β-D-maltopyranoside (DDM) (Anatrace, Inc. Maumee, OH), EDTA-free complete protease inhibitor cocktail (Roche, Basel, Switzerland), and incubated at 4°C for one hour. After centrifugation at 27,000 ×g for 30 min, the supernatant was loaded onto a strep-tactin (IBA Lifesciences, Göttingen, Germany) column equilibrated with the lysis buffer. The column was washed with 50 column volumes of Buffer A and 0.3% DDM, followed by additional washes with 50 column volumes of Buffer A and 0.1% DDM, and with 50 column volumes of Buffer A and 0.02% DDM. The S protein was eluted by Buffer A containing 0.02% DDM and 5 mM desthiobiotin (IBA Lifesciences). The protein was further purified by gel filtration chromatography on a Superose 6 10/300 column (GE Healthcare, Chicago, IL) in a buffer containing 25 mM Tris-HCl, pH 7.5, 150 mM NaCl, 0.02% DDM.

Another stable cell line was also generated for large-scale production of monomeric soluble ACE2 protein. Purification of the ACE2 protein was carried out as previously described (*59*). Briefly, the stably transfected cells were incubated in Expi293 expression medium containing 1% Pen Strep and 1.0 μg/ml puromycin to a density of ~4-5×10^6^/mL, the cell supernatant was harvested by centrifugation at 3,000 xg for 30 min and loaded onto a column packed with Ni Sepharose excel resin (Cytiva Life Sciences, Marlborough, MA). The column was washed with a buffer containing 20 mM Tris-HCl, pH 7.5, 300 mM NaCl and 10 mM imidazole. The ACE2 protein was eluted with a buffer containing 300 mM imidazole, and further purified by gel filtration chromatography on HiLoad 16/600 Superdex 200 pg column (GE Healthcare) in 25 mM Tris-HCl, pH 7.5 and 150 mM NaCl.

To produce the linear MSP2N2 protein, the expression construct was transformed into *E. coli* BL21 Star (DE3) (Thermo Fisher Scientific), and the cells were grown at 37 °C in LB medium with 50 μg/mL kanamycin to an OD_600_ of 0.8 and then induced with 1 mM isopropyl-β-d-thiogalactopyranoside (IPTG; Sigma-Aldrich, St. Louis, MO) at 30 °C for 5 h, and harvested by centrifugation. Purification and circularization of the lsMSP2N2 protein were carried out as previously described (*42*, *61*). Briefly, the cells were completely resuspended in a buffer (40 mM Tris-HCl, pH 8.0, 300 mM NaCl) and lysed by sonication. The cell lysate was clarified by centrifugation at 30,000 ×g for 30 min and the supernatant was loaded onto a column packed with Ni-NTA agarose (Qiagen, Hilden, Germany). The column was sequentially washed with buffer I (40 mM Tris-HCl, pH 8.0, 300 mM NaCl, 1% Trion X-100), buffer II (40 mM Tris-HCl, pH 8.0, 300 mM NaCl, 50 mM sodium cholate, 20 mM imidazole), and buffer III (40 mM Tris-HCl, pH 8.0, 300 mM NaCl, 50 mM imidazole). The lsMSP2N2 protein was eluted with 40 mM Tris-HCl, pH 8.0, 300 mM NaCl, 400 mM imidazole. To remove the N-terminal His-tag, the protein was treated with TEV protease in the TEV buffer (20 mM Tris-HCl, pH 8.0, 100 mM NaCl, 1 mM EDTA, 1 mM DTT) at room temperature for 3 hrs. To produce circularized protein - csMSP2N2 while removing its C-terminal His-tag, the sortase eSrt was mixed with the TEV-treated lsMSP2N2 with a molar ratio of 1:20 in a circularization buffer (30 mM Tris-HCl, pH 7.5, 150 mM NaCl, 10 mM CaCl_2_, 1 mM DTT), and incubated at 37 °C for 3 hrs. The circularization reaction was passed through a Ni-NTA agarose column to remove the histagged protein and collect the flow-through, which contained the csMSP2N2 protein. The protein was further purified by anion exchange chromatography on a HiTrap Q HP column (Cytiva Life Sciences), dialyzed against 20 mM Tris-HCl, pH 7.5 and 100 mM NaCl, flash frozen in liquid nitrogen and stored at −80 °C.

The trimeric ACE2 protein and S-specific monoclonal antibodies were described previously (*50*, *59*)

#### Reconstitution of the spike protein in nanodiscs

To reconstitute the spike protein in nanodiscs, soy extract polar lipid (Avanti, Birmingham, AL) in chloroform was first dried under a nitrogen stream and dried further overnight in a vacuum desiccator. Lipid films were dissolved in a buffer containing 25 mM Tris 7.5, 150 mM NaCl, 2% DDM, 0.4% CHS, vortexed vigorously and sonicated until the solution became clear. The S protein and lipids were mixed and incubated on ice for 30 min. The csMSP2N2 was added to the mixture with a spike:csMSP2N2:lipid molar ratio of 1:8:700 and incubated on ice for another 30 min. Biobeads SM2 (Bio-Rad, Hercules, CA) were added to remove detergents from the mixture and initiate the reconstitution. After gentle rotation at 4 °C overnight, Bio-beads were removed through filtration. To induce the conformational transition of the S protein from the prefusion state to the postfusion state in the nanodiscs, the reconsitution mixture was added with soluble ACE2 protein at 6 μM and incubated at room temperature for 30 min. A control experiment confirmed the prefusion S protein was stable at room temperature in absence of ACE2. The ACE2-induced postfusion S-nanodisc sample was purified by gel filtration chromatography on Superose 6 Increase 10/300 column (GE Healthcare) in a buffer containing 25 mM Tris-HCl, pH 7.5, 150 mM NaCl. Peak fractions containing the postfusion S-nanodisc sample were used for cryo-EM analysis.

#### Negative stain EM

To prepare grids, 4 μl of freshly purified prefusion or postfusion S nanodisc sample was adsorbed to a glow-discharged carbon-coated copper grid (Electron Microscopy Sciences, Hatfield, PA), washed with deionized water, and stained with freshly prepared 1.5% uranyl formate. Images were recorded at room temperature on the Phillips CM10 transmission electron microscope with a nominal magnification of 52,000×. Particles were auto-picked and 2D class averages were generated using RELION 4.0.0 (*62*).

#### Cryo-EM sample preparation and data collection

To prepare cryo EM grids, the postfusion S-nanodisc sample at 1.5 mg/ml was applied to a 1.2/1.3 Quantifoil gold grid (Quantifoil Micro Tools GmbH), which had been glow discharged with a PELCO easiGlow™ Glow Discharge Cleaning system (Ted Pella, Inc.) for 60 s at 15 mA. Grids were immediately plunge-frozen in liquid ethane using a Vitrobot Mark IV (ThermoFisher Scientific), and excess protein was blotted away using grade 595 filter paper (Ted Pella, Inc.) with a blotting time of 4 s, a blotting force of −12 at 4°C with 100% humidity. The grids were screened for ice thickness and particle distribution. Selected grids were used to acquire images with a Titan Krios transmission electron microscope (ThermoFisher Scientific) operated at 300 keV and equipped with a BioQuantum GIF/K3 direct electron detector. Automated data collection was carried out using SerialEM version 4.0.5 (*63*) at a nominal magnification of 105,000× and the K3 detector in counting mode (calibrated pixel size, 0.825 Å) at an exposure rate of ~13.8 electrons per pixel per second. Each movie adds a total accumulated electron exposure of ~50.6/51.3 e-/Å^2^, fractionated in 50/51 frames. Data sets were acquired using a defocus range of 0.8-2.2 μm.

#### Image processing and 3D reconstructions

All data were processed using cryoSPARC v.3.3.1 (*43*) and RELION. Drift correction for cryo-EM images was performed using patch mode, and contrast transfer function (CTF) estimated by patch mode. Motion corrected sums with dose-weighting were used for all other image processing. Blob picking was performed for particle picking. 18,339,830 particles were extracted from 142,99 images using a box size of 600 Å (downsizing to 128Å) from data set I, and 57,656,504 particles from 17,028 images from data set II. The two sets of particles were processed separately and each subjected to 8-10 rounds of 2D classification, giving 1,532,458 and 3,885,698 good particles, respectively. An initial model was produced using ab-initio reconstruction in cryoSPARC based on selected 2D class averages from data set I.

For data set I, good particles from 2D classification were used for one round of heterogeneous classification with six copies of the initial model as the reference in C1 symmetry. Three major classes with total 57.6% of the particles showing clear structural features were subjected to two additional rounds of heterogeneous refinement with six copies of the initial model as the reference in C1 symmetry. The major class with a nanodisc shape at one end was re-extracted to an unbinned box size of 600Å and subjected to one round of 2D classification to remove bad particles, followed by one round of non-uniform refinement in C1 symmetry to produce a map at 3.18Å resolution from 153,658 particles. Another round of non-uniform refinement in C3 symmetry was performed and improved the resolution to 3.0Å. To further improve the local resolution in the nanodisc, these particles were transferred to RELION and subjected to one round of focus-classification with a mask covering the entire protein, leading to a major class (78.9% particles) with continuous density at the transmembrane region. This class with 123,298 particles was imported back to cryoSPARC, and subjected to another round of non-uniform refinement in C3 symmetry, giving a map at 2.96Å resolution. One round of particle-subtraction was performed to remove noises from the nanodiscs, followed by one round of local refinement with an overall mask in C3 symmetry to produce a map at 2.84Å resolution. Another round of local refinement in C3 symmetry focused on the transmembrane region was performed subsequently to improve its resolution, giving a map at 3.35Å resolution. These overall and locally refined maps showed clear density for transmembrane helices, but the resolution was not high enough for unambiguously modeling the structure. Data set II was therefore collected with the same batch of the S-nanodisc sample.

For data set II, the selected particles from 2D classification were used for one round of heterogeneous classification with six copies of the initial model as the reference in C1 symmetry. Two major classes with total 37.8% of the particles showing clear structural features were subjected to two additional rounds of heterogeneous refinement with six copies of the initial model as the reference in C1 symmetry. The major class was re-extracted to an unbinned box size of 600Å and subjected to one round of 2D classification, followed by one round of duplicates removing, non-uniform refinement in C1 symmetry, giving a map at 3.4Å resolution from 131,604 particles. These particles were combined with the particle set from data set I after the focus-classification, and subjected to one round of non-uniform refinement in C3 symmetry, CTF refinement, and another round of non-uniform refinement in C3 symmetry, giving a final overall map at 2.86Å resolution from the combined 254,902 particles. Particle-subtraction was also performed to reduce noises from the nanodiscs, followed by two rounds of local refinement with a different mask in size each time, giving a map at 3.26Å resolution in the transmembrane region with showing clear sidechain density for the transmembrane helices. The best maps from data set I and the combined data set were all used for model building.

All resolutions were reported from the gold-standard Fourier shell correlation (FSC) using the 0.143 criterion. Density maps were corrected from the modulation transfer function of the K3 detector and sharpened by applying a temperature factor that was estimated using sharpening tools in cryoSPARC. Local resolution was also determined using cryoSPARC.

#### Model building

The initial template for model building was our postfusion S structure (PDB ID: 7XRA). Several rounds of manual building were performed in Coot. The model around the transmembrane region was first refined in Phenix (*64*) against the locally refined maps at 3.26/3.35Å resolution, and further refined in Phenix against the 2.9Å overall map. Iteratively, refinement was performed in both Phenix (real space refinement) and ISOLDE (*65*), and the Phenix refinement strategy included minimization_global, local_grid_search, and adp, with rotamer, Ramachandran, and reference-model restraints, using 7KRA as the reference models. The local map from data set I had stronger density for the CT region and was used for modeling the CT; the map from the combined data set map had better sidechain density for the transmembrane helices and thus used for modeling them. The refinement statistics are summarized in Table S1.

We used AlphaFold2 (*53*) implementation in the ColabFold (*66*) notebooks running on Google Colaboratory and our new postfusion S2 structure as a template to predict the structures of the S2 trimers from other SARS-CoV-2 variants. Default settings were used with Amber relaxation; the sequences were entered in tandem and separated by a semicolon for prediction of a trimer. AlphaFold was performed once with each of the trained models, which were verified for consistency, and the best model was chosen based on pLDDT (predicted Local Distance Difference Test) score. Structural biology applications used in this project were compiled and configured by SBGrid (*67*).

#### Western blot

Full-length S protein samples were resolved in 4-15% Mini-Protean TGX gel (Bio-Rad) and transferred onto PVDF membranes (Millipore, Billerica, MA) by an Iblot2 (Invitrogen by Thermo Fisher Scientific). Membranes were blocked with 5% skimmed milk in PBS for 1 hour and incubated with a SARS-CoV-2 (2019-nCoV) Spike RBD Antibody (Sino Biological Inc., Beijing, China, Cat: 40592-T62) for another hour at room temperature. Alkaline phosphatase conjugated anti-Rabbit IgG (1:5000) (Sigma-Aldrich, St. Louis, MO) was used as a secondary antibody. Proteins were visualized using one-step NBT/BCIP substrates (Promega, Madison, WI).

#### Flow cytometry

Expi293F cells were grown in Expi293 expression medium. Cell surface display DNA constructs for the SARS-CoV-2 G614 or its mutants or S2 together with a plasmid expressing blue fluorescent protein (BFP) were transiently transfected into Expi293F cells using ExpiFectamine 293 reagent (ThermoFisher Scientific) per manufacturer’s instruction. Two days after transfection, the cells were stained with primary antibodies at 5 μg/ml concentration. An Alexa Fluor 647 conjugated donkey anti-human IgG Fc F(ab’)2 fragment (Jackson ImmunoResearch, West Grove, PA) was used as secondary antibody at the concentration of 5 μg/ml. Cells were run through an Intellicyt iQue Screener Plus flow cytometer. Cells gated for positive BFP expression were analyzed for antibody binding.

#### Cell-cell fusion assay

The cell-cell fusion assay, based on the α-complementation of *E. coli* β-galactosidase, was conducted to quantify the fusion activity mediated by SARS-CoV-2 S protein, as described previously (*20*). Briefly, the full-length G614 S or its mutants (10 μg) and the α fragment of *E. coli* β-galactosidase construct (10 μg), or the full-length ACE2 construct (5 μg) together with the ω fragment of *E. coli* β-galactosidase construct (10 μg), were transfected in HEK 293T cells using polyethylenimine (PEI) (80 μg). After incubation at 37°C for 5 hrs, the medium was aspirated and replaced with complete DMEM (1% Pen Strep, 1% GlutaMax and 10% FBS), followed by incubation at 37°C for additional 19 hrs. The cells were detached using PBS and resuspended in complete DMEM. 50 μl S-expressing cells (1.0×10^6^ cells/ml) were mixed with 50 μl ACE2-expressing cells (1.0×10^6^ cells/ml) to allow cell-cell fusion to proceed at 37°C for 4 hrs. Cell-cell fusion activity was quantified using a chemiluminescent assay system, Gal-Screen (Applied Biosystems, Foster City, CA), following the standard protocol recommended by the manufacturer. The substrate was added to the mixture of the cells and allowed to react for 90 min in dark at room temperature. The luminescence signal was recorded with a Synergy Neo plate reader (Biotek, Winooski, VT).

## Acknowledgments

We thank the SBGrid team for computing support, computing resources from S. Harrison, and S. Harrison for critical reading of the manuscript. We acknowledge support for COVID-19 related structural biology research at Harvard from the Nancy Lurie Marks Family Foundation and the Massachusetts Consortium on Pathogen Readiness (MassCPR). This work was supported by COVID-19 Awards by MassCPR (to B.C.), Fast grant by Emergent Ventures (to B.C.), and NIH grants AI127193 (to B.C. and James Chou), AI147884 (to B.C.) and AI141002 (to B.C.).

## Author Contribution

B.C., J.Z. and W.S. conceived the project. W.S. produced the full-length S protein and reconstituted in nanodiscs with help from Y.C. and H.P.. Y.C. also created the new MSP construct and established initial conditions for nanodisc reconstitution. W.S. and J.Z. prepared cryo grids and performed EM data collection with contributions from M.L.M., K.S. and C.X.. J.Z. processed the cryo-EM data, built and refined the atomic models. H.C. and J.L. created the mutant constructs. H.Z. performed the flow cytometry experiment. J.Z. and W.S. designed the mutant constructs; W.S. and J.Z. carried out the cell-cell fusion assay. J.V. and S.R.V. contributed to plasmid preparation, cell culture and protein production. All authors analyzed the data. B.C., J.Z. and W.S. wrote the manuscript with input from all other authors.

## Data Availability

The atomic structure coordinates and EM maps are deposited in the EMDataBank under the accession number: xxx. All other related data generated during and/or analyzed during the current study are available from the corresponding authors on reasonable request.

## Reference

1. R. P. Rand, V. A. Parsegian, Physical force considerations in model and biological membranes. Can J Biochem Cell Biol 62, 752–759 (1984).

2. V. A. Parsegian, N. Fuller, R. P. Rand, Measured work of deformation and repulsion of lecithin bilayers. Proc Natl Acad Sci U S A 76, 2750–2754 (1979).

3. S. C. Harrison, Viral membrane fusion. Virology 479-480, 498–507 (2015).

4. M. Kielian, Mechanisms of Virus Membrane Fusion Proteins. Annu Rev Virol 1, 171–189 (2014).

5. W. Weissenhorn et al., Structural basis for membrane fusion by enveloped viruses. Mol Membr Biol 16, 3–9 (1999).

6. C. B. Jackson, M. Farzan, B. Chen, H. Choe, Mechanisms of SARS-CoV-2 entry into cells. Nat Rev Mol Cell Biol 23, 3–20 (2022).

7. M. A. Tortorici, D. Veesler, Structural insights into coronavirus entry. Adv Virus Res 105, 93–116 (2019).

8. M. Hoffmann et al., SARS-CoV-2 Cell Entry Depends on ACE2 and TMPRSS2 and Is Blocked by a Clinically Proven Protease Inhibitor. Cell 181, 271–280 e278 (2020).

9. M. M. Zhao et al., Cathepsin L plays a key role in SARS-CoV-2 infection in humans and humanized mice and is a promising target for new drug development. Signal Transduct Target Ther 6, 134 (2021).

10. W. Weissenhorn, A. Dessen, S. C. Harrison, J. J. Skehel, D. C. Wiley, Atomic structure of the ectodomain from HIV-1 gp41. Nature 387, 426–430 (1997).

11. D. C. Chan, D. Fass, J. M. Berger, P. S. Kim, Core structure of gp41 from the HIV envelope glycoprotein. Cell 89, 263–273 (1997).

12. F. Wu et al., A new coronavirus associated with human respiratory disease in China. Nature 579, 265–269 (2020).

13. D. Wrapp et al., Cryo-EM structure of the 2019-nCoV spike in the prefusion conformation. Science 367, 1260–1263 (2020).

14. A. C. Walls et al., Structure, function and antigenicity of the SARS-CoV-2 spike glycoprotein. Cell, DOI: 10.1016/j.cell.2020.1002.1058 (2020).

15. J. Lan et al., Structure of the SARS-CoV-2 spike receptor-binding domain bound to the ACE2 receptor. Nature 581, 215–220 (2020).

16. R. Yan et al., Structural basis for the recognition of SARS-CoV-2 by full-length human ACE2. Science 367, 1444–1448 (2020).

17. J. Shang et al., Structural basis of receptor recognition by SARS-CoV-2. Nature 581, 221–224 (2020).

18. Q. Wang et al., Structural and Functional Basis of SARS-CoV-2 Entry by Using Human ACE2. Cell 181, 894–904 e899 (2020).

19. S. Xia et al., Inhibition of SARS-CoV-2 (previously 2019-nCoV) infection by a highly potent pan-coronavirus fusion inhibitor targeting its spike protein that harbors a high capacity to mediate membrane fusion. Cell Res 30, 343–355 (2020).

20. Y. Cai et al., Distinct conformational states of SARS-CoV-2 spike protein. Science 369, 1586–1592 (2020).

21. S. Bangaru et al., Structural analysis of full-length SARS-CoV-2 spike protein from an advanced vaccine candidate. Science 370, 1089–1094 (2020).

22. B. Turonova et al., In situ structural analysis of SARS-CoV-2 spike reveals flexibility mediated by three hinges. Science 370, 203–208 (2020).

23. Z. Ke et al., Structures and distributions of SARS-CoV-2 spike proteins on intact virions. Nature 588, 498–502 (2020).

24. H. Yao et al., Molecular Architecture of the SARS-CoV-2 Virus. Cell 183, 730–738 e713 (2020).

25. C. Liu et al., The Architecture of Inactivated SARS-CoV-2 with Postfusion Spikes Revealed by Cryo-EM and Cryo-ET. Structure 28, 1218–1224.e1214 (2020).

26. B. Schroth-Diez et al., The role of the transmembrane and of the intraviral domain of glycoproteins in membrane fusion of enveloped viruses. Biosci Rep 20, 571–595 (2000).

27. B. J. Bosch, C. A. de Haan, S. L. Smits, P. J. Rottier, Spike protein assembly into the coronavirion: exploring the limits of its sequence requirements. Virology 334, 306–318 (2005).

28. E. Lontok, E. Corse, C. E. Machamer, Intracellular targeting signals contribute to localization of coronavirus spike proteins near the virus assembly site. J Virol 78, 5913–5922 (2004).

29. C. M. Petit et al., Genetic analysis of the SARS-coronavirus spike glycoprotein functional domains involved in cell-surface expression and cell-to-cell fusion. Virology 341, 215–230 (2005).

30. R. Ye, C. Montalto-Morrison, P. S. Masters, Genetic analysis of determinants for spike glycoprotein assembly into murine coronavirus virions: distinct roles for charge-rich and cysteine-rich regions of the endodomain. J Virol 78, 9904–9917 (2004).

31. S. Belouzard, J. K. Millet, B. N. Licitra, G. R. Whittaker, Mechanisms of coronavirus cell entry mediated by the viral spike protein. Viruses 4, 1011–1033 (2012).

32. B. Sainz, Jr., J. M. Rausch, W. R. Gallaher, R. F. Garry, W. C. Wimley, Identification and characterization of the putative fusion peptide of the severe acute respiratory syndrome-associated coronavirus spike protein. J Virol 79, 7195–7206 (2005).

33. L. G. Basso, E. F. Vicente, E. Crusca, Jr., E. M. Cilli, A. J. Costa-Filho, SARS-CoV fusion peptides induce membrane surface ordering and curvature. Sci Rep 6, 37131 (2016).

34. I. G. Madu, S. L. Roth, S. Belouzard, G. R. Whittaker, Characterization of a highly conserved domain within the severe acute respiratory syndrome coronavirus spike protein S2 domain with characteristics of a viral fusion peptide. J Virol 83, 7411–7421 (2009).

35. A. L. Lai, J. K. Millet, S. Daniel, J. H. Freed, G. R. Whittaker, The SARS-CoV Fusion Peptide Forms an Extended Bipartite Fusion Platform that Perturbs Membrane Order in a Calcium-Dependent Manner. J Mol Biol 429, 3875–3892 (2017).

36. B. J. Bosch et al., Severe acute respiratory syndrome coronavirus (SARS-CoV) infection inhibition using spike protein heptad repeat-derived peptides. Proc Natl Acad Sci U S A 101, 8455–8460 (2004).

37. C. Dacon et al., Broadly neutralizing antibodies target the coronavirus fusion peptide. Science 377, 728–735 (2022).

38. J. S. Low et al., ACE2-binding exposes the SARS-CoV-2 fusion peptide to broadly neutralizing coronavirus antibodies. Science 377, 735–742 (2022).

39. J. Guillen, A. J. Perez-Berna, M. R. Moreno, J. Villalain, Identification of the membrane-active regions of the severe acute respiratory syndrome coronavirus spike membrane glycoprotein using a 16/18-mer peptide scan: implications for the viral fusion mechanism. J Virol 79, 1743–1752 (2005).

40. B. Sainz, Jr., J. M. Rausch, W. R. Gallaher, R. F. Garry, W. C. Wimley, The aromatic domain of the coronavirus class I viral fusion protein induces membrane permeabilization: putative role during viral entry. Biochemistry 44, 947–958 (2005).

41. Y. V. Grinkova, I. G. Denisov, S. G. Sligar, Engineering extended membrane scaffold proteins for self-assembly of soluble nanoscale lipid bilayers. Protein Eng Des Sel 23, 843–848 (2010).

42. N. T. Johansen et al., Circularized and solubility-enhanced MSPs facilitate simple and high-yield production of stable nanodiscs for studies of membrane proteins in solution. FEBS J 286, 1734–1751 (2019).

43. A. Punjani, J. L. Rubinstein, D. J. Fleet, M. A. Brubaker, cryoSPARC: algorithms for rapid unsupervised cryo-EM structure determination. Nat Methods 14, 290–296 (2017).

44. X. Fan, D. Cao, L. Kong, X. Zhang, Cryo-EM analysis of the post-fusion structure of the SARS-CoV spike glycoprotein. Nat Commun 11, 3618 (2020).

45. A. A. Ramadan et al., Identification of SARS-CoV-2 Spike Palmitoylation Inhibitors That Results in Release of Attenuated Virus with Reduced Infectivity. Viruses 14, (2022).

46. C. F. Tien et al., Glycosylation and S-palmitoylation regulate SARS-CoV-2 spike protein intracellular trafficking. iScience 25, 104709 (2022).

47. Y. Modis, S. Ogata, D. Clements, S. C. Harrison, Structure of the dengue virus envelope protein after membrane fusion. Nature 427, 313–319 (2004).

48. D. A. Steinhauer, S. A. Wharton, J. J. Skehel, D. C. Wiley, Studies of the membrane fusion activities of fusion peptide mutants of influenza virus hemagglutinin. J Virol 69, 6643–6651 (1995).

49. I. A. Wilson, J. J. Skehel, D. C. Wiley, Structure of the haemagglutinin membrane glycoprotein of influenza virus at 3 A resolution. Nature 289, 366–373 (1981).

50. P. Tong et al., Memory B cell repertoire for recognition of evolving SARS-CoV-2 spike. Cell 184, 4969–4980 e4915 (2021).

51. A. Baum et al., Antibody cocktail to SARS-CoV-2 spike protein prevents rapid mutational escape seen with individual antibodies. Science 369, 1014–1018 (2020).

52. X. Chi et al., A neutralizing human antibody binds to the N-terminal domain of the Spike protein of SARS-CoV-2. Science 369, 650–655 (2020).

53. J. Jumper et al., Highly accurate protein structure prediction with AlphaFold. Nature 596, 583–589 (2021).

54. A. J. B. Kreutzberger et al., SARS-CoV-2 requires acidic pH to infect cells. Proc Natl Acad Sci U S A 119, e2209514119 (2022).

55. J. L. Lorieau, J. M. Louis, A. Bax, The complete influenza hemagglutinin fusion domain adopts a tight helical hairpin arrangement at the lipid:water interface. Proc Natl Acad Sci U S A 107, 11341–11346 (2010).

56. J. Corver, R. Broer, P. van Kasteren, W. Spaan, Mutagenesis of the transmembrane domain of the SARS coronavirus spike glycoprotein: refinement of the requirements for SARS coronavirus cell entry. Virol J 6, 230 (2009).

57. Q. Fu, J. J. Chou, A Trimeric Hydrophobic Zipper Mediates the Intramembrane Assembly of SARS-CoV-2 Spike. J Am Chem Soc 143, 8543–8546 (2021).

58. J. Zhang et al., Structural impact on SARS-CoV-2 spike protein by D614G substitution. Science 372, 525–530 (2021).

59. T. Xiao et al., A trimeric human angiotensin-converting enzyme 2 as an anti-SARS-CoV-2 agent. Nat Struct Mol Biol 28, 202–209 (2021).

60. J. Chen et al., HIV-1 ENVELOPE. Effect of the cytoplasmic domain on antigenic characteristics of HIV-1 envelope glycoprotein. Science 349, 191–195 (2015).

61. T. K. Ritchie et al., Chapter 11 - Reconstitution of membrane proteins in phospholipid bilayer nanodiscs. Methods Enzymol 464, 211–231 (2009).

62. S. H. Scheres, RELION: implementation of a Bayesian approach to cryo-EM structure determination. J Struct Biol 180, 519–530 (2012).

63. D. N. Mastronarde, Automated electron microscope tomography using robust prediction of specimen movements. J Struct Biol 152, 36–51 (2005).

64. P. D. Adams et al., PHENIX: a comprehensive Python-based system for macromolecular structure solution. Acta Crystallogr D Biol Crystallogr 66, 213–221 (2010).

65. T. I. Croll, ISOLDE: a physically realistic environment for model building into low-resolution electron-density maps. Acta Crystallogr D Struct Biol 74, 519–530 (2018).

66. M. Mirdita et al., ColabFold: making protein folding accessible to all. Nat Methods 19, 679–682 (2022).

67. A. Morin et al., Collaboration gets the most out of software. Elife 2, e01456 (2013).

68. F. Madeira et al., The EMBL-EBI search and sequence analysis tools APIs in 2019. Nucleic Acids Res 47, W636–W641 (2019).

69. X. Robert, P. Gouet, Deciphering key features in protein structures with the new ENDscript server. Nucleic Acids Res 42, W320–324 (2014).

