## Supplementary materials for "Cryo-EM structure of SARS-CoV-2 postfusion spike in membrane"

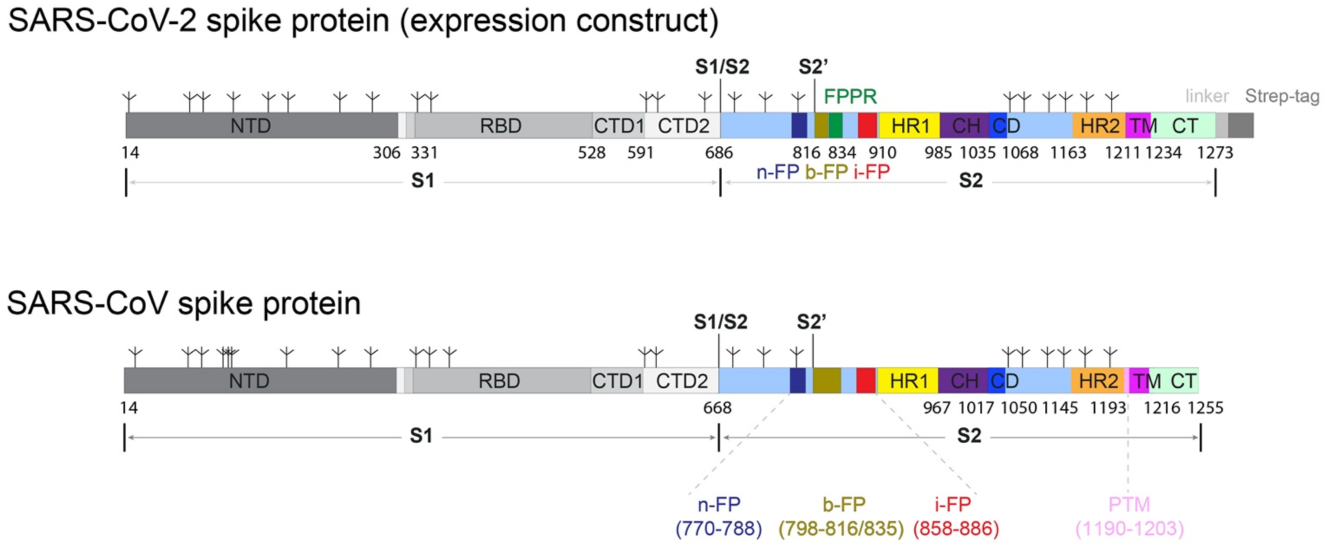


**Figure S1. Schematic representations of the full-length SARS-CoV-2 and SARS-CoV spike proteins.** Segments of S1 and S2 include: NTD, N-terminal domain; RBD, receptor-binding domain; CTD1, C-terminal domain 1; CTD2, C-terminal domain 2; S1/S2, the furin cleavage site at the S1/S2 boundary; S2', S2' cleavage site; FP, fusion peptide; FPPR, fusion peptide proximal region; HR1, heptad repeat 1; CH, central helix region; CD, connector domain; HR2, heptad repeat 2; TM, transmembrane segment; CT, cytoplasmic tail; and tree-like symbols for glycans. Based on studies of the SARS-CoV S protein, three membranotropic regions in the SARS-CoV S2 have been suggested as putative fusion peptides, including a segment of residues 770-788, named the N-terminal FP (n-FP); a segment of residues 798-816 or 798-835, widely accepted as the "bona fide" FP (b-FP); and a segment of residues 858-886, also known as the internal FP (i-FP). A segment of residues 1190-1203 is called the pretransmembrane domain (pre-TM).

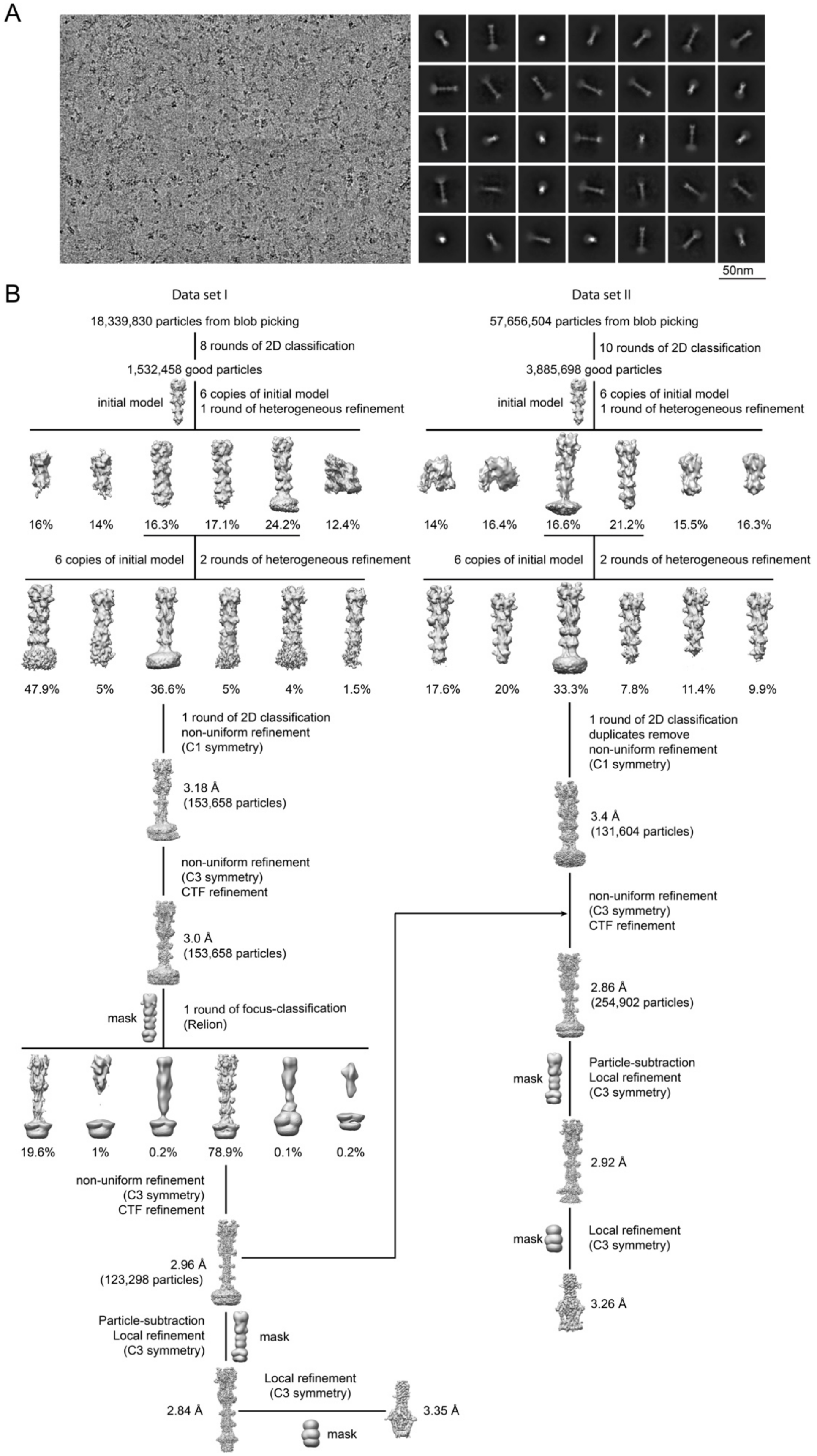

A
50nm
B
Data set I
18,339,830 particles from blob picking
8 rounds of 2D classification
1,532,458 good particles
initial model
6 copies of initial model
1 round of heterogeneous refinement
16%
14%
16.3%
17.1%
24.2%
12.4%
6 copies of initial model
2 rounds of heterogeneous refinement
47.9%
5%
36.6%
5%
4%
1.5%
1 round of 2D classification
non-uniform refinement
(C1 symmetry)
3.18 Å
(153,658 particles)
non-uniform refinement
(C3 symmetry)
CTF refinement
3.0 Å
(153,658 particles)
mask
1 round of focus-classification
(Relion)
19.6%
1%
0.2%
78.9%
0.1%
0.2%
non-uniform refinement
(C3 symmetry)
CTF refinement
2.96 Å
(123,298 particles)
Particle-subtraction
Local refinement
(C3 symmetry)
mask
2.84 Å
Local refinement
(C3 symmetry)
mask
3.35 Å
Data set II
57,656,504 particles from blob picking
10 rounds of 2D classification
3,885,698 good particles
initial model
6 copies of initial model
1 round of heterogeneous refinement
14%
16.4%
16.6%
21.2%
15.5%
16.3%
6 copies of initial model
2 rounds of heterogeneous refinement
17.6%
20%
33.3%
7.8%
11.4%
9.9%
1 round of 2D classification
duplicates remove
non-uniform refinement
(C1 symmetry)
3.4 Å
(131,604 particles)
non-uniform refinement
(C3 symmetry)
CTF refinement
2.86 Å
(254,902 particles)
mask
Particle-subtraction
Local refinement
(C3 symmetry)
2.92 Å
mask
Local refinement
(C3 symmetry)
3.26 Å

**Figure S2. Cryo-EM analysis of the postfusion S2 trimer.** Top, representative micrograph, and 2D averages (box dimension: 500Å) of the cryo-EM particle images of the postfusion S2 trimer in nanodiscs. Bottom, data processing workflow for structure determination.

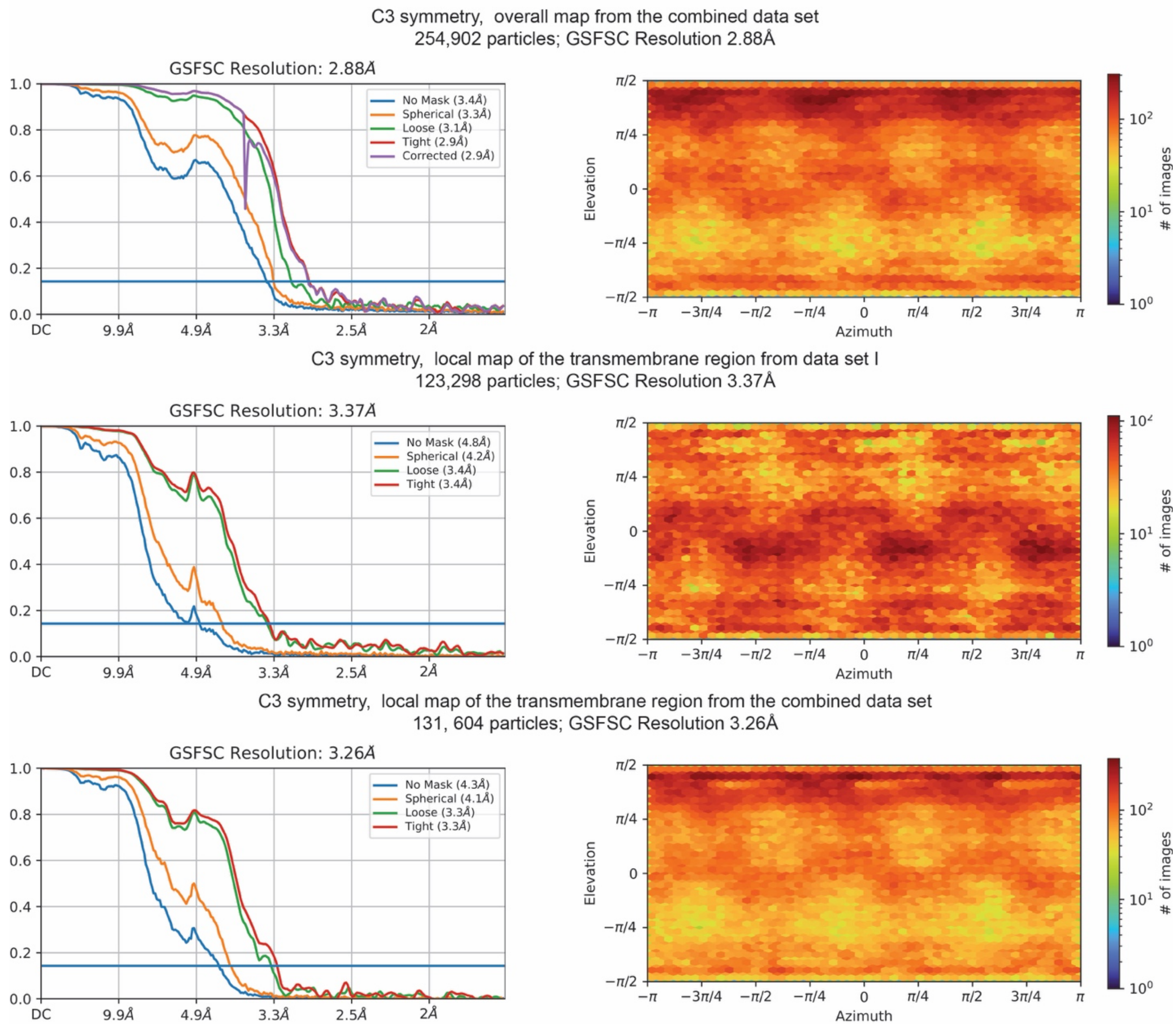


**Figure S3. Analysis of the postfusion S2 trimer structures by cryoSPARC.** Gold standard FSC curves of the three refined 3D reconstructions of the postfusion S2 trimer and the corresponding cryoSPARC output for particle distribution of each reconstruction.

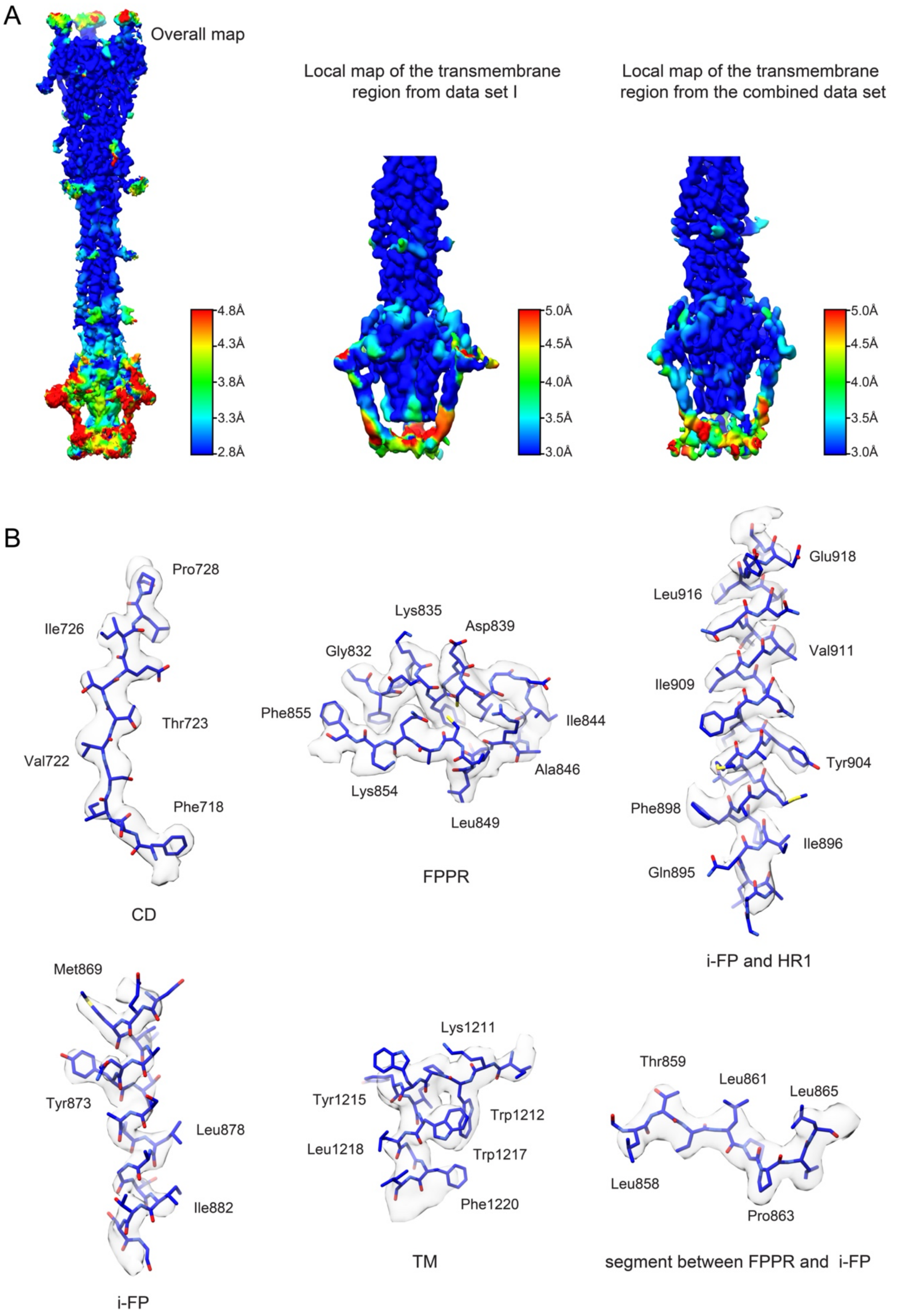
A
Overall map
Local map of the transmembrane region from data set I
Local map of the transmembrane region from the combined data set
4.8Å
4.3Å
3.8Å
3.3Å
2.8Å
5.0Å
4.5Å
4.0Å
3.5Å
3.0Å
5.0Å
4.5Å
4.0Å
3.5Å
3.0Å
B
Pro728
Ile726
Thr723
Val722
Phe718
CD
Lys835
Asp839
Gly832
Phe855
Ile844
Ala846
Lys854
Leu849
FPPR
Glu918
Leu916
Val911
Ile909
Tyr904
Phe898
Ile896
Gln895
i-FP and HR1
Met869
Tyr873
Leu878
Ile882
i-FP
Lys1211
Tyr1215
Trp1212
Leu1218
Trp1217
Phe1220
TM
Thr859
Leu861
Leu865
Leu858
Pro863
segment between FPPR and i-FP

**Figure S4. Additional analysis of the postfusion S2 trimer structure.** (**A**) 3D reconstructions of the postfusion S2 trimer from the overall refinement and local refinement are colored according to local resolution estimated by ResMap with the FSC=0.143 criterion. (**B**) Representative density in gray surface representation from the EM maps of the postfusion S2.

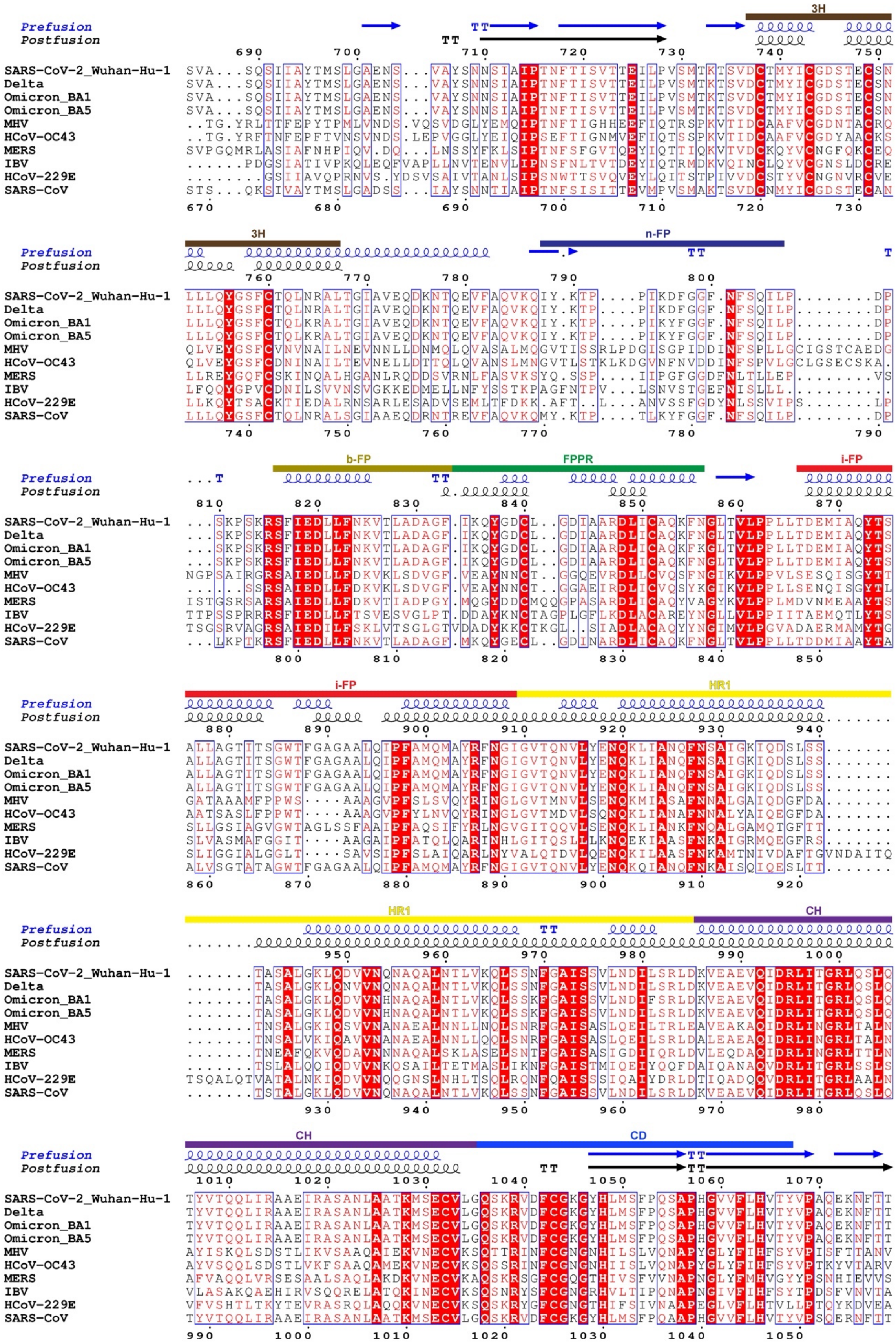
3H
Prefusion
Postfusion
SARS-CoV-2_Wuhan-Hu-1 SVA...SQSIIAYTMSLGAENS...VAYSNNSIAIPTNFTISVTTEILPVSMTKTSVDCTMYICGDSTECSN
Delta SVA...SQSIIAYTMSLGAENS...VAYSNNSIAIPTNFTISVTTEILPVSMTKTSVDCTMYICGDSTECSN
Omicron_BA1 SVA...SQSIIAYTMSLGAENS...VAYSNNSIAIPTNFTISVTTEILPVSMTKTSVDCTMYICGDSTECSN
Omicron_BA5 SVA...SQSIIAYTMSLGAENS...VAYSNNSIAIPTNFTISVTTEILPVSMTKTSVDCTMYICGDSTECSN
MHV ..TG.YRLTTFEPYTPMLVNDS.VQSVDGLYEMQIPTNFTIGHHEEFIQTRSPKVTIDCAAFVCGDNTACRQ
HCoV-OC43 ..TG.YRFTNFEPFTVNSVNDS.LEPVGGLYEIQIPSEFTIGNMVEFIQTSSPKVTIDCAAFVCGDYAACKS
MERS SVPGQMRLASIAFNHPIQV.DQ...LNSSYFKLSIPTNFSFGVTQEYIQTTIQKVTVDCKQYVCNGFQKCEQ
IBV ......PDGSIATIVPKQLEQFVAPLLNVTENVLIPNSFNLTVTDEYIQTRMDKVQINCLQYVCGNSLDCRE
HCoV-229E ........GSIIAVQPRNVS.YDSVSAIVTANLSIPSNWTTSVQVEYLQITSTPIVVDCSTYVCNGNVRCVE
SARS-CoV STS...QKSIVAYTMSLGADSS...IAYSNNTIAIPTNFSISITTEVMPVSMAKTSVDCNMYICGDSTECAN
3H
n-FP
Prefusion
Postfusion
SARS-CoV-2_Wuhan-Hu-1 LLLQYGSFCTQLNRALTGIAVEQDKNTQEVFAQVKQIY.KTP....PIKDFGGF.NFSQILP........DP
Delta LLLQYGSFCTQLNRALTGIAVEQDKNTQEVFAQVKQIY.KTP....PIKDFGGF.NFSQILP........DP
Omicron_BA1 LLLQYGSFCTQLKRALTGIAVEQDKNTQEVFAQVKQIY.KTP....PIKYFGGF.NFSQILP........DP
Omicron_BA5 LLLQYGSFCTQLKRALTGIAVEQDKNTQEVFAQVKQIY.KTP....PIKYFGGF.NFSQILP........DP
MHV QLVEYGSFCVNVNAILNEVNNLLDNMQLQVASALMQGVTISSRLPDGISGPIDDINFSPLLGCIGSTCAEDG
HCoV-OC43 QLVEYGSFCDNINAILTEVNELLDTTQLQVANSLMNGVTLSTKLKDGVNFNVDDINFSPVLGCLGSECSKA.
MERS LLREYGQFCSKINQALHGANLRQDDSVRNLFASVKSYQ.SSP....IIPGFGGDFNLTLLEP........VS
IBV LFQQYGPVCDNILSVVNSVGKKEDMELLNFYSSTKPAGFNTPV...LSNVSTGEFNISLLL..........
HCoV-229E LLKQYTSACKTIEDALRNSARLESADVSEMLTFDKK.AFT..L...ANVSSFGDYNLSSVIPS.......LP
SARS-CoV LLLQYGSFCTQLNRALSGIAAEQDRNTREVFAQVKQMY.KTP....TLKYFGGF.NFSQILP........DP
b-FP
FPPR
i-FP
Prefusion
Postfusion
SARS-CoV-2_Wuhan-Hu-1 ...SKPSKRSFIEDLLFNKVTLADAGF.IKQYGDCL..GDIAARDLICAQKFNGLTVLPPLLTDEMIAQYTS
Delta ...SKPSKRSFIEDLLFNKVTLADAGF.IKQYGDCL..GDIAARDLICAQKFNGLTVLPPLLTDEMIAQYTS
Omicron_BA1 ...SKPSKRSFIEDLLFNKVTLADAGF.IKQYGDCL..GDIAARDLICAQKFKGLTVLPPLLTDEMIAQYTS
Omicron_BA5 ...SKPSKRSFIEDLLFNKVTLADAGF.IKQYGDCL..GDIAARDLICAQKFNGLTVLPPLLTDEMIAQYTS
MHV NGPSAIRGRSAIEDLLFDKVKLSDVGF.VEAYNNCT..GGQEVRDLLCVQSFNGIKVLPPVLSESQISGYTT
HCoV-OC43 ......SSRSAIEDLLFDKVKLSDVGF.VEAYNNCT..GGAEIRDLICVQSYKGIKVLPPLLSENQISGYTL
MERS ISTGSRSARSAIEDLLFDKVTIADPGY.MQGYDDCMQQGPASARDLICAQYVAGYKVLPPLMDVNMEAAYTS
IBV TTPSSPRRRSFIEDLLFTSVESVGLPT.DDAYKNCTAGPLGFLKDLACAREYNGLLVLPPIITAEMQTLYTS
HCoV-229E TSGSRVAGRSAIEDILFSKLVTSGLGTVDADYKKCTKGL..SIADLACAQYYNGIMVLPGVADAERMAMYTG
SARS-CoV ...LKPTKRSFIEDLLFNKVTLADAGF.MKQYGECL..GDINARDLICAQKFNGLTVLPPLLTDDMIAAYTA
i-FP
HR1
Prefusion
Postfusion
SARS-CoV-2_Wuhan-Hu-1 ALLAGTITSGWTFGAGAALQIPFAMQMAYRFNGIGVTQNVLYENQKLIANQFNSAIGKIQDSLSS.......
Delta ALLAGTITSGWTFGAGAALQIPFAMQMAYRFNGIGVTQNVLYENQKLIANQFNSAIGKIQDSLSS.......
Omicron_BA1 ALLAGTITSGWTFGAGAALQIPFAMQMAYRFNGIGVTQNVLYENQKLIANQFNSAIGKIQDSLSS.......
Omicron_BA5 ALLAGTITSGWTFGAGAALQIPFAMQMAYRFNGIGVTQNVLYENQKLIANQFNSAIGKIQDSLSS.......
MHV GATAAAMFPPWS....AAAGVPFSLSVQYRINGLGVTMNVLSENQKMIASAFNNALGAIQDGFDA.......
HCoV-OC43 AATSASLFPPWT....AAAGVPFYLNVQYRINGLGVTMDVLSQNQKLIANAFNNALYAIQEGFDA.......
MERS SLLGSIAGVGWTAGLSSFAAIPFAQSIFYRLNGVGITQQVLSENQKLIANKFNQALGAMQTGFTT.......
IBV SLVASMAFGGIT....AAGAIPFATQLQARINHLGITQSLLLKNQEKIAASFNKAIGRMQEGFRS.......
HCoV-229E SLIGGIALGGLT....SAVSIPFSLAIQARLNYVALQTDVLQENQKILAASFNKAMTNIVDAFTGVNDAITQ
SARS-CoV ALVSGTATAGWTFGAGAALQIPFAMQMAYRFNGIGVTQNVLYENQKQIANQFNKAISQIQESLTT.......
HR1
CH
Prefusion
Postfusion
SARS-CoV-2_Wuhan-Hu-1 .......TASALGKLQDVVNQNAQALNTLVKQLSSNFGAISSVLNDILSRLDKVEAEVQIDRLITGRLQSLQ
Delta .......TASALGKLQNVVNQNAQALNTLVKQLSSNFGAISSVLNDILSRLDKVEAEVQIDRLITGRLQSLQ
Omicron_BA1 .......TASALGKLQDVVNHNAQALNTLVKQLSSKFGAISSVLNDIFSRLDKVEAEVQIDRLITGRLQSLQ
Omicron_BA5 .......TASALGKLQDVVNHNAQALNTLVKQLSSKFGAISSVLNDILSRLDKVEAEVQIDRLITGRLQSLQ
MHV .......TNSALGKIQSVVNANAEALNNLLNQLSNRFGAISASLQEILTRLEAVEAKAQIDRLINGRLTALN
HCoV-OC43 .......TNSALVKIQAVVNANAEALNNLLQQLSNRFGAISASLQEILSRLDALEAEAQIDRLINGRLTALN
MERS .......TNEAFQKVQDAVNNNAQALSKLASELSNTFGAISASIGDIIQRLDVLEQDAQIDRLINGRLTTLN
IBV .......TSLALQQIQDVVNKQSAILTETMASLIKNFGAISTMIQEIYQQFDAIQANAQVDRLITGRLSSLS
HCoV-229E TSQALQTVATALNKIQDVVNQQGNSLNHLTSQLRQNFQAISSSIQAIYDRLDTIQADQQVDRLITGRLAALN
SARS-CoV .......TSTALGKLQDVVNQNAQALNTLVKQLSSNFGAISSVLNDILSRLDKVEAEVQIDRLITGRLQSLQ
CH
CD
Prefusion
Postfusion
SARS-CoV-2_Wuhan-Hu-1 TYVTQQLIRAAEIRASANLAATKMSECVLGQSKRVDFCGKGYHLMSFPQSAPHGVVFLHVTYVPAQEKNFTT
Delta TYVTQQLIRAAEIRASANLAATKMSECVLGQSKRVDFCGKGYHLMSFPQSAPHGVVFLHVTYVPAQEKNFTT
Omicron_BA1 TYVTQQLIRAAEIRASANLAATKMSECVLGQSKRVDFCGKGYHLMSFPQSAPHGVVFLHVTYVPAQEKNFTT
Omicron_BA5 TYVTQQLIRAAEIRASANLAATKMSECVLGQSKRVDFCGKGYHLMSFPQSAPHGVVFLHVTYVPAQEKNFTT
MHV AYISKQLSDSTLIKVSAAQAIEKVNECVKSQTTRINFCGNGNHILSLVQNAPYGLYFIHFSYVPISFTTANV
HCoV-OC43 AYVSQQLSDSTLVKFSAAQAMEKVNECVKSQSSRINFCGNGNHIISLVQNAPYGLYFIHFSYVPTKYVTARV
MERS AFVAQQLVRSESAALSAQLAKDKVNECVKAQSKRSGFCGQGTHIVSFVVNAPNGLYFMHVGYYPSNHIEVVS
IBV VLASAKQAEHIRVSQQRELATQKINECVKSQSNRYSFCGNGRHVLTIPQNAPNGIVFIHFSYTPDSFVNVTA
HCoV-229E VFVSHTLTKYTEVRASRQLAQQKVNECVKSQSKRYGFCGNGTHIFSIVNAAPEGLVFLHTVLLPTQYKDVEA
SARS-CoV TYVTQQLIRAAEIRASANLAATKMSECVLGQSKRVDFCGKGYHLMSFPQAAPHGVVFLHVTYVPSQERNFTT

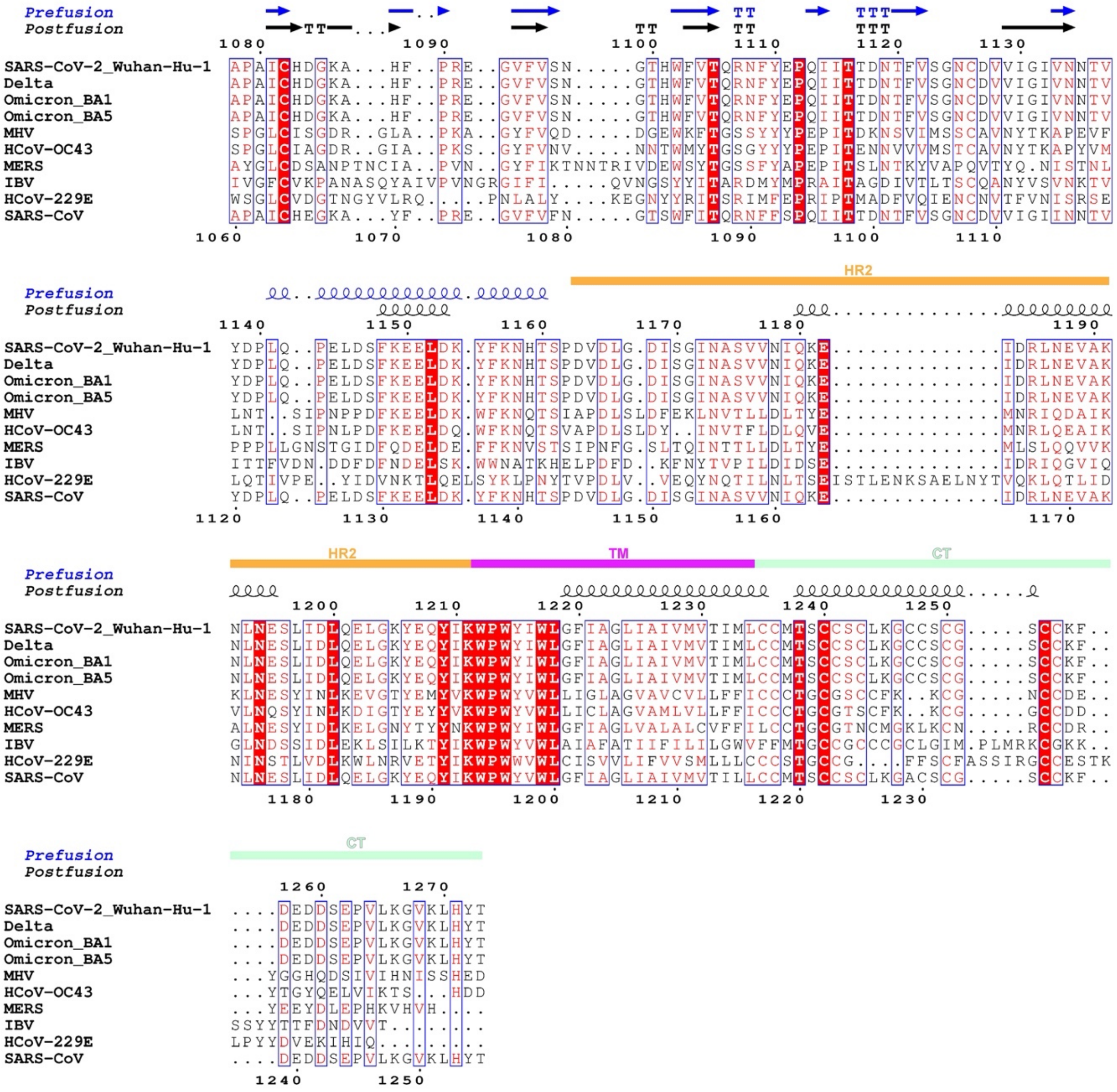

**Figure S5. Sequence alignment of the S2 proteins from various coronaviruses**. The intact S2 proteins from various coronaviruses are aligned by Clustal Omega (*68*) and displayed using the online server ESPript 3.0 (*69*), with highly conserved residues highlighted in red. The SARS-CoV-2 and SARS-CoV spikes are numbered, and various segments color-coded with rectangle bars. The secondary structures of various segments in both the prefusion and postfusion conformations are indicated.

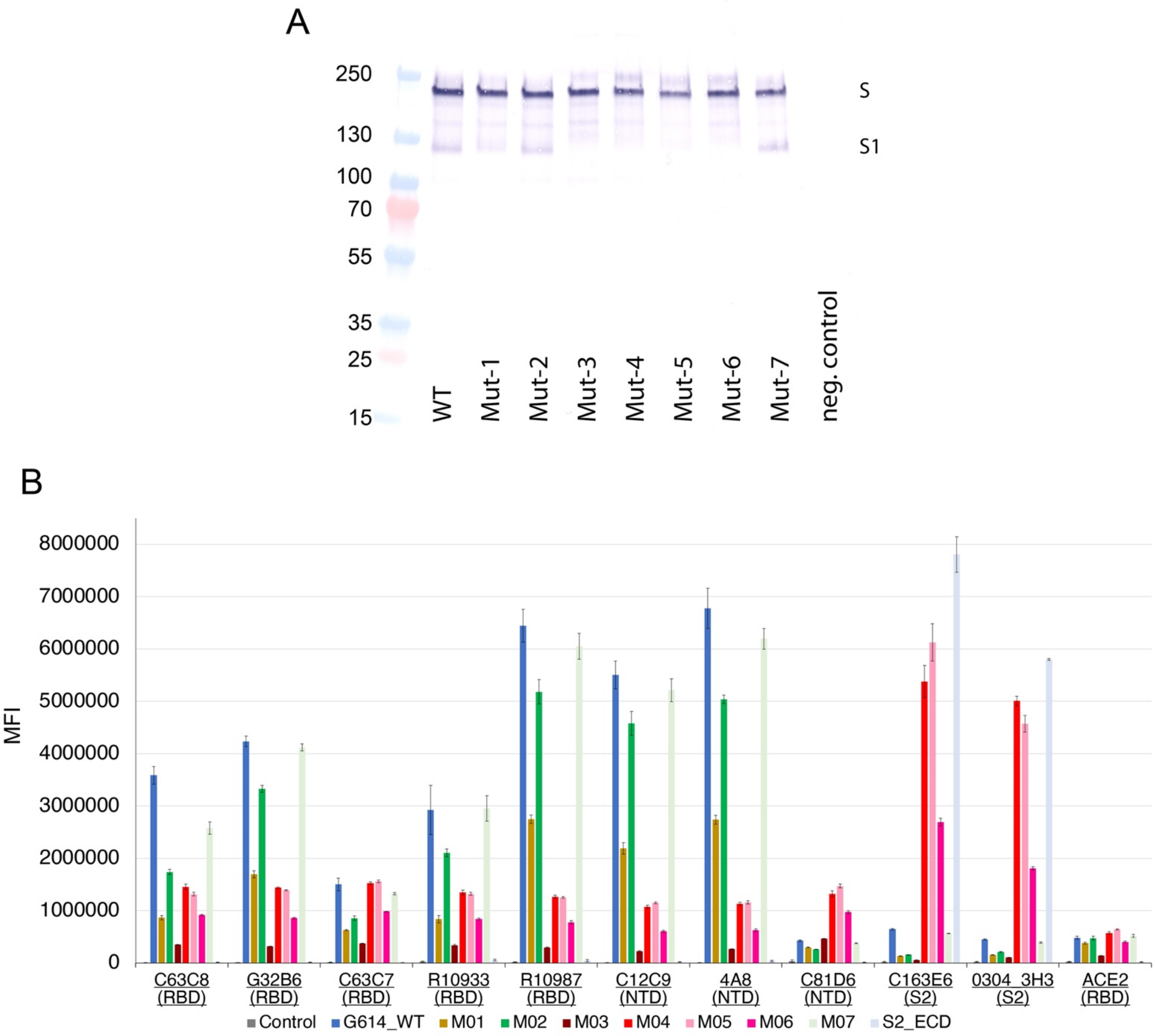


**Figure S6. Expression and antigenic properties of the mutant spike proteins assessed by flow cytometry.** (**A**) Expression and processing of the full-length G614 S construct and its mutants in HEK293 cells. S protein samples prepared from HEK293 cells transiently transfected with 10 μg of the full-length S expression plasmids were detected by anti-RBD polyclonal antibodies. Bands for the uncleaved S and S1 fragment are indicated. The experiment was repeated four times with similar results. (**B**) Antibody binding to the full-length G614 S protein and its mutants, as well as an S2 construct expressed on the cell surfaces analyzed by flow cytometry. The antibodies and their targets are indicated. A designed ACE2-based fusion inhibitor ACE2615-foldon-T27W was used for detecting receptor binding (*59*). MFI, mean fluorescent intensity. The error bars represent standard errors of mean from measurements using three independently transfected cell samples. The flow cytometry assays were repeated three times with essentially identical results.

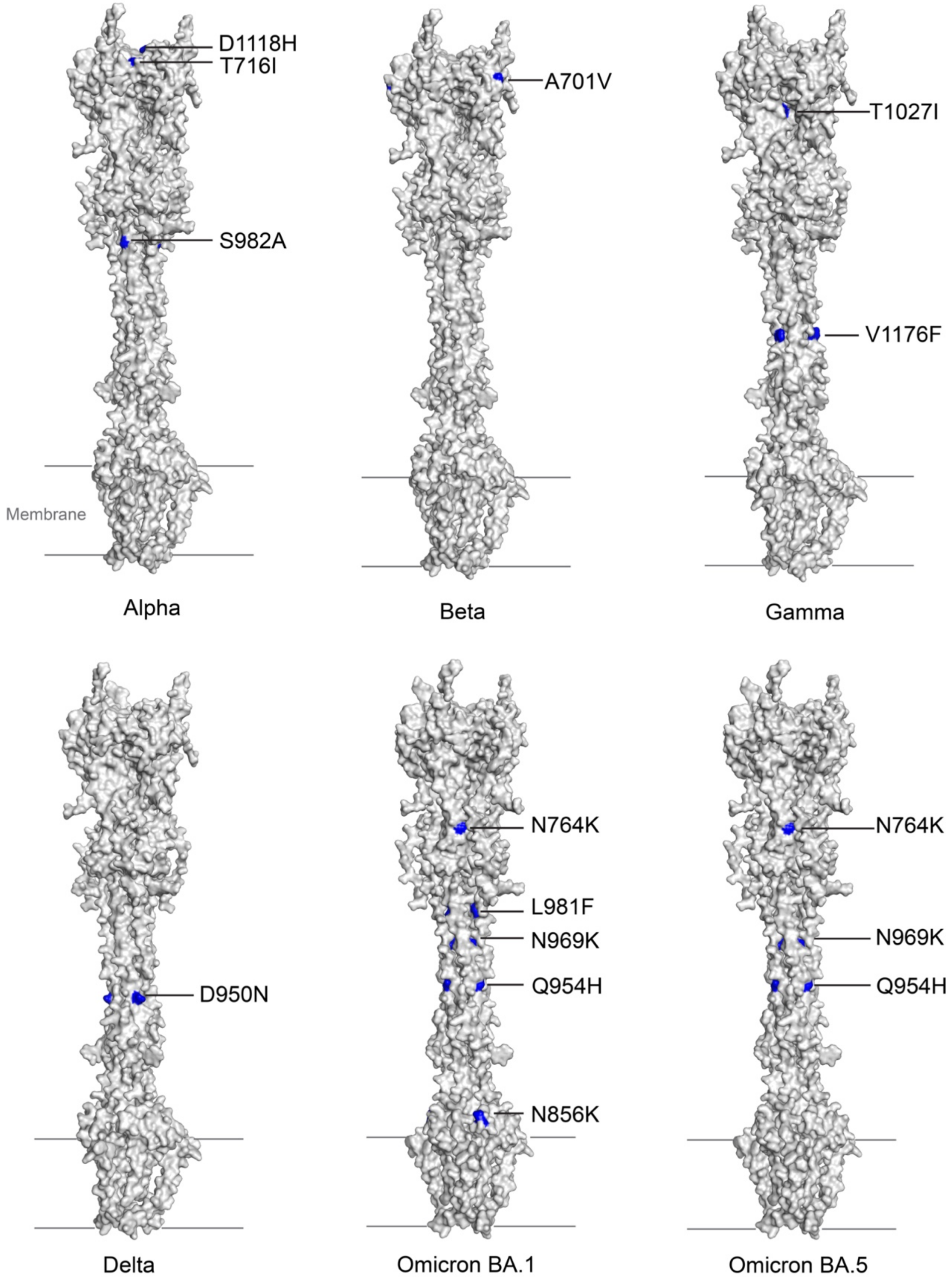


**Figure S7. Modeling of the S2 trimer structures of SARS-CoV-2 variants by AlphaFold.** The models of the postfusion S trimers of Alpha, Beta, Gamma, Delta, Omicron BA.1 and Omicron BA.5 variants of concern, generated by AlphaFold are shown in the surface representation in gray with the mutations in each variant highlighted in blue.

### Table S1. Cryo-EM statistics.

| **EM data collection and reconstruction statistics** | | |
|---|---|---|
| Protein | SARS-CoV-2 postfusion spike protein in nanodiscs | |
| Microscope | Titan Krios | |
| Voltage(kV) | 300 | |
| Detector | Gatan K3 | |
| Magnification(nominal) | 105,000 | |
| Energy filter slit width (eV) | 20 | |
| Calibrated pixel size (Å/pix) | 0.825 | |
| Exposure rate (e$^-$/pix/sec) | 13.8 | 13.884 |
| Frames per exposure | 50 | 51 |
| Total electron exposure (e$^-$/Å$^2$) | 50.6 | 51.2624 |
| Exposure per frame (e$^-$/Å$^2$) | 1.01 | 1.01 |
| Defocus range (μm) | -1.2, -2.2 | -0.8,-2.2 |
| Automation software | SerialEM | |
| # of Micrographs used | 142,99 | 17,028 |
| Particles extracted | 18,339,830 | 57,656,504 |
| Particles after 2D classification | 1,532,458 | 3,885,698 |

| Map | Overall map | Local map from data set I | Local map from the combined data set |
|---|---|---|---|
| Total # of refined particles | 254,902 | 123,298 | 254,902 |
| Symmetry imposed | C3 | C3 | C3 |
| Map sharpening B-factor | -93.1 | -112.3 | -122.5 |
| Map resolution (Å) | 2.9 | 3.4 | 3.3 |
| FSC threshold(Å) | 0.143 | 0.143 | 0.143 |

| Model refinement and validation statistics | |
|---|---|
| PDB | |
| Composition | |
| Amino acids | 1527 |
| Glycans | 39 |
| RMSD bonds (Å) | 0.014 |
| RMSD angles (◦) | 1.927 |
| Mean B-factors | |
| Amino acids | 75 |
| Glycans | 99 |
| Ramachandran | |
| Favored (%) | 92.81 |
| Allowed(%) | 5.54 |
| Outliers(%) | 1.65 |
| Rotamer outliers (%) | 2.76 |
| Clash score | 1.13 |
| C-beta outliers (%) | 0.63 |
| CaBLAM outliers (%) | 3.73 |
| CC (mask) | 0.63 |
| CC (volume) | 0.63 |
| MolProbity score | 1.61 |
| EMRinger score | 1.34 |
